# Mechanical Tension Actively Triggers RhoA-Mediated Cell Extrusion

**DOI:** 10.64898/2026.07.08.737158

**Authors:** Fanny Wodrascka, Tianxiang Ma, Pablo Gottheil, Robin Durand, Lucas Anger, Andreas Schoenit, Mallica Pandya, Manon Arnaud, Tien Dang, Siavash Monfared, Guillaume Charras, René-Marc Mège, Amin Doostmohammadi, Benoit Ladoux, Simon de Beco

## Abstract

Cell extrusion is a fundamental process in tissue homeostasis, morphogenesis, and cancer progression, facilitating the removal of cells either alive or through apoptosis. While biochemical signaling pathways are known to regulate extrusion, recent advances have underscored the importance of mechanical forces in this process. Here, using optogenetic control of RhoA activation in epithelial monolayers combined with Bayesian Inversion Stress Microscopy (BISM) and three-dimensional cell-based modeling, we uncover a counterintuitive mechanism whereby elevated tension, instead of stabilizing the monolayer, actively drives extrusion in highly contractile cells. We show that local RhoA activation enhances myosin II–dependent contractility and F-actin reorganization, which promotes cell stiffening, resulting in localized tension buildup. The ensuing tensile stress amplifies vertical mechanical fluctuations, which are tightly associated with cell extrusion. Remarkably, these tension-induced extrusions occur both apically and basally. Furthermore, our findings show that RhoA-mediated contractility is not merely an effector of extrusion but also an active promoter of basal extrusion, independently of caspase activation. Our study demonstrates that tensile stress can directly initiate extrusion events and bias their outcome toward apical or basal fates. By identifying tension as a driver rather than a suppressor of extrusion, this work revises current models of epithelial homeostasis and highlights mechanical control as a targetable axis in disease and regeneration.

## Introduction

Epithelial tissues maintain their structure and function through a precise balance between cell proliferation and elimination^1^. This balance generates a characteristic “homeostatic pressure,” reflecting the equilibrium between cell division and loss. When it is disrupted, overcrowding or tissue degeneration can occur^2^. A key process in regulating this pressure is cell extrusion, a conserved mechanism by which cells are expelled from the monolayer, either alive or as dead cells during apoptosis^3^, epithelial-mesenchymal transition (EMT)^4^, or precancerous cell invasion^5^. Cell signaling pathways have long been recognized as critical regulators of cell extrusion. Apoptotic extrusion is tightly coordinated with a cascade of biochemical cues, involving the release of Sphingosine-1-Phosphate to the neighboring cells resulting in RhoA activation to form a contractile actomyosin ring necessary for the apoptotic cell to be expelled from the tissue^2,6^. RhoA signaling plays a crucial role by activating myosin-II contraction in the extruding cell and i neighboring cells at opposing cell-cell junctions. This coordinated action ensures the maintenance of tissue integrity by preventing gaps between cells. Furthermore, contractile apoptotic cells communicate with their neighbors through E-cadherin-mediated mechanotransduction involving Myosin VI as well as p114 and p115 RhoGEF^7^. Conversely, RhoA inhibition along neighboring junctions releases tension and permits junctional elongation as the delaminated cell shrinks^8^.

Accordingly, mechanical forces have emerged as major regulators of cell extrusion, through tissue compaction, cell crowding^2,9^, and compressive stresses surrounding extrusion sites^10,11^. Additionally, compressive stresses influence the fate of delaminated cell that can undergo apoptosis^2,9,10^ or remain alive^11^. In turn, the contractile forces exerted by apoptotic cells participate in shaping the surrounding tissue during embryonic development^12,13^. Importantly, dysregulation of cell elimination or defects in extruded cell fate can contribute to diseases, including cancer^14,15^. Understanding how cells sense, process and respond to mechanical inputs is therefore critical for uncovering the mechanisms governing extrusion and apoptosis. Mechanistically, cell extrusion involves cellular mechanical responses underscored by cytoskeletal remodeling in the extruded cell and its neighbors^16–18^. This is exemplified by the formation of actin cables in dying cells and the involvement of cryptic lamellipodia or purse-string-like mechanisms for cell removal^7,8,19–21^. This intricate coordination of cellular contractile activities suggests that cell extrusion is not a purely cell-autonomous process but is instead influenced by the local mechanical environment.

These observations prompted us to ask whether localized versus global RhoA activity differentially regulates extrusion. We hypothesized that fluctuations of cellular contractility would be sufficient to trigger this process.

Using optogenetics to control RhoA semi-quantitatively^22–24^, we investigated how RhoA-induced contractility at the multicellular scale generates tensile stress and influences both cell extrusion and the fate of extruding cells. Our observations demonstrate that contractility-induced tension can promote cell extrusion. When cells were cultured on thick collagen gels, these extrusions were detected in both apical and basal directions, with basal extrusions occurring independently of caspase activation. To elucidate the underlying mechanisms, we integrated experimental biophysical techniques—including Traction Force Microscopy (TFM)^25^, Bayesian Inversion Stress Microscopy (BISM)^26^, and indentation-based stiffness measurements—with in silico three-dimensional phase field modeling. Our results show that RhoA activation enhances myosin II–dependent contractility, leading to local cell stiffening via F-actin remodeling^27,28^. The resulting mechanical mismatch between RhoA-activated and neighboring non-activated cells drives asymmetric transmission of active stress, producing localized tension within the activated region. This elevated tension, in turn, amplifies vertical (out-of-plane) mechanical fluctuations, which are associated with basal or apical extrusion. Together, these findings identify RhoA not only as an effector but also as a promoter of cell elimination, revealing a tension-driven, non-apoptotic route to extrusion that expands our understanding of how mechanical forces govern epithelial tissue dynamics.

## Results

### Optogenetic activation of RhoA generates tensile stress in epithelial monolayers

We used optogenetics to control RhoA activity and cell contractility with high spatiotemporal resolution in Madin-Darby Canine Kidney (MDCK) epithelial cells. Expression of CIBN-GFP-CAAX and ArhGEF11-CRY2-mCherry (MDCK Opto-RhoA) allows the relocalization of ArhGEF11, a specific GEF of RhoA, from the cytoplasm to the plasma membrane upon blue light illumination^22,23^ (Figure 1 A,B, Video S1), resulting in RhoA activation and increased contractility. MDCK Opto-RhoA or Opto-Ctrl cells (lacking ArhGEF11 fusion to CRY2-mCherry) were grown on 15 kPa PolyDiMethylSiloxane (PDMS) surfaces coated with fluorescent beads in order to measure traction forces^29^ (Figure 1C). Illumination of MDCK Opto-RhoA cells within 150 µm × 150 µm square regions triggered a rapid, localized, and sustained increase in traction forces (Figure 1 D,E), whereas no change was observed in MDCK Opto-Ctrl cells. It is worth noting that MDCK Opto-RhoA monolayers, even in the absence of illumination, exhibited elevated baseline traction forces, likely reflecting leakiness in the optogenetic system as previously observed^23^. Illuminated Opto-RhoA cells exhibited a greater number of focal adhesions and elevated levels of p-MLC2 (Figure S1 A,B), consistent with increased contractility.

**Figure 1:**
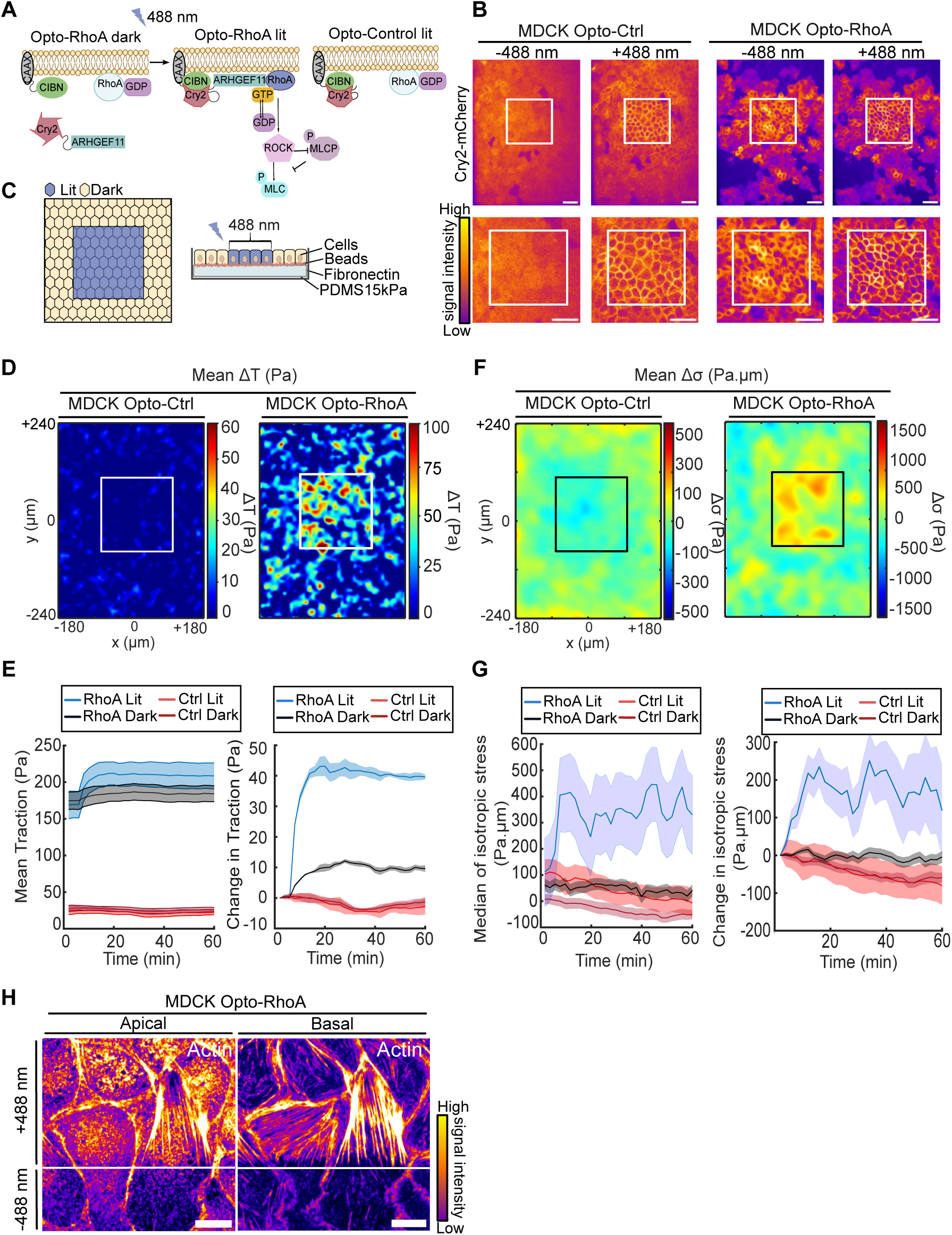
Control of tensile stress in space and time through RhoA opto-activation. **A** Schematic of the optogenetic system used to trigger RhoA activation in MDCK cells. (left) CRY2-ArhGEF11-mCherry is recruited to the plasma membrane following blue light-stimulated binding to CIBN, leading to RhoA activation and subsequent myosin contraction. (Right) ARHGEF11 is not fused to CRY2 in the control cell line (Opto-Ctrl). **B** Examples of membrane recruitment of CRY2-mCherry or ArhGEF11-CRY2-mCherry following activation at 488 nm in MDCK Opto-Ctrl (left) and Opto-RhoA cells (right) respectively, within a 150 x 150 µm blue-light illuminated square (white outline). Bottom: inset magnification. Scale bar: 50 µm. **C** Experimental setup representation of localized blue optogenetic stimulation combined with Traction Force Microscopy (TFM) experiment: MDCK Opto-RhoA cells are seeded on 15 kPa PDMS coated with fluorescent beads and fibronectin and illuminated in a restricted area using a FRAP module. **D, F** Maps of mean change in traction forces **(D)** and isotropic stress **(F)**, 60 minutes after light activation in the 150 µm x 150 µm square (white outline), in MDCK Opto-Ctrl (left, N=2 n=30), or Opto-RhoA cells (right, N=2 n=38). **E,G** Quantification of absolute values (left) and change (right) in traction forces (**E**, N=3 n=36) and isotropic stress (**G**, N=2 n=38) in illuminated regions for Opto-RhoA (blue) and Opto-Ctrl (red) or surrounding unstimulated regions (black and dark red, respectively). Data are mean +/- S.E.M. **H** Representative images of Opto-RhoA cells activated in a square and stained for actin. Display of apical view (left) and basal view (right) show the enrichment and remodeling of actin restricted to the stimulated area. The continuous line represents the border between the stimulated (+488 nm) and unstimulated (−488 nm) areas. Scale bar: 10 µm. See also Figure S1, S2 and Video S1.

Analysis of p-MLC2 distribution revealed a similar basal enrichment in Opto-Ctrl and Opto-RhoA monolayers, with no significant difference in the basal-to-apical fluorescence ratio (Figure S1 C,D), indicating that optogenetic RhoA activation increases contractility without disrupting its polarized organization. However, Bayesian Inversion Stress Microscopy (BISM, see Methods)^26,30^ demonstrated that RhoA-induced contractility reshapes the stress landscape, resulting in elevated isotropic tensile stress within the illuminated region (Figure 1 F,G). This finding was confirmed by laser ablation, which revealed a faster and more pronounced recoil in illuminated regions of MDCK Opto-RhoA monolayers compared to non-illuminated areas (Figure S1 E,F). This rise in tensile stress in the illuminated regions coincided with a marked reduction in cell velocity and mean-square displacement (MSD) (Figure S1 G,H). Given the mechanosensitive nature of cadherin-based junctions^31^, we next investigated whether the increase in tensile stress affected cell-cell junctions. We found reduced levels of E-cadherin and α-catenin in Opto-RhoA monolayers (Figure S1B), consistent with previously described mechanosensitive regulation of E-cadherin dynamics^32,33^.

To investigate the mechanical origin of this tension buildup in silico, we developed a minimal three-dimensional (3D) model of an epithelial monolayer, incorporating both passive and active mechanics of deformable cells (see Supplementary Methods for details and parametrization). The model employs a multiphase field framework^34,35^, and has been validated to accurately capture both in-plane force transmission and out-of-plane stress components responsible for cell extrusion^18,36^. Briefly, each deformable cell is described by a phase-field variable whose dynamics are governed by energy minimization and force balance. Passive contributions include cell-cell and cell-substrate adhesive and repulsive interactions, as well as the cellular mechanical properties including elasticity and compressibility. Active contributions include intercellular stresses that mimic force transmission through cadherin-based junctions^37^, as well as self-propulsion forces oriented by contact inhibition of locomotion^38^.

To make the model directly comparable to our experimental system, we incorporated experimentally measured changes in cell mechanical properties induced by RhoA activation. Previous studies have shown that myosin-driven contractility is closely linked to cell stiffening^39^, primarily through reorganization of F-actin^40^. Consistently, we observed pronounced F-actin accumulation in the RhoA-activated regions (Figure 1H), and indentation experiments confirmed significantly increased stiffness (Figure S2A). We asked whether this contractility-associated stiffness change could account for the tension buildup. To this end, we represented this mechanical change by increasing γ in the activated region, which modulates interfacial mechanics and thereby the effective deformability of the simulated cells^41,42^ (see Supplementary Methods for details, and Figure S2B), while keeping all other parameters unchanged. Under these conditions, a robust and spatially confined rise in in-plane tension emerged in the activated region (Figure S2 C,D), closely matching the profile of stress observed experimentally (Figure 1F). This model configuration also recapitulated the reduced cell velocity and MSD in the activated regions (Figure S2 E,F), as well as reduced intercellular adhesion (Figure S3), qualitatively consistent with the experimentally observed decrease in E-cadherin (Figure S1B).

To further validate this model, we tested two alternative modifications associated with RhoA activation — increased contractile active stress along basal actomyosin fibers (see Supplementary Methods for details) and increased cell–substrate adhesion, corresponding to the elevation in phosphorylated myosin levels and greater number of focal adhesions observed in illuminated Opto-RhoA cells — neither of which reproduced the experimentally observed tension buildup (Figure S4). We therefore incorporated stiffness as a consequence of RhoA activation in our model and computed the simulated in-plane force density across the monolayer to identify the physical origin of tension buildup. At the interface between activated and non-activated regions, the total force (defined here as the sum of all simulated in-plane force contributions, including passive cell-cell and cell-substrate adhesive/repulsive forces, active intercellular forces, and self-propulsion forces) pointed outward from the activated region (Figure S2G, black arrow). By pulling the boundary of the activated region away from its center, this outward force provides a stretching contribution, consistent with the accumulation of tensile stress within the activated region. To identify the origin of this stretching force, we separately examined the active and passive mechanical components. As established (see Supplemental Information), active intercellular stress is proportional to the cell deformation tensor. Due to stiffness mismatch, cells in the activated region are rounder (low deformation), while surrounding cells are more elongated (high deformation), creating a gradient in the deformation and in turn, an active intercellular force contribution pointing outward (Figure S2G, blue arrows). In contrast, the passive force acting on the activated region at the interfaces arises from inward-directed cell–cell repulsion and outward-directed cell-cell adhesion; under our parameter regime, their net effect points inward, imposing a compressive constraint on the activated region (Figure S2G, gray arrows). In addition to these, we found that the self-propulsion force pointed outward at the interface between activated and non-activated regions (Figure S2G, red arrow, see SI Note). Thus, tension accumulation arises chiefly from active intercellular and self-propulsion forces created by stiffness asymmetries, with passive forces acting as a confining counterbalance.

Altogether, our results showed that optogenetic activation of RhoA allowed for a fast and local increase of cell contractility, highlighted by the assembly of actin stress fibers and increased stiffness, which ultimately leads to the accumulation of tensile isotropic intercellular stress within the activated region (Figure S2H).

### RhoA-induced tension triggers cell extrusion

We then asked whether tensile stress, generated by RhoA activation, could also affect extrusion, by quantifying the extrusion rates of Opto-RhoA and Opto-Ctrl cells in illuminated and non-illuminated regions (Figure 2A, Video S2). RhoA activation increased the extrusion rate by ∼1.7-fold (Figure 2B). This was confirmed by stimulation of MDCK Opto-Ctrl cells with the RhoA activator CN03 which also increased the cell extrusion rate (Figure S5). In fact, even non-illuminated Opto-RhoA cells show higher extrusion rates than Opto-Ctrl ones, which are less contractile (Figure S5). Instead, inhibition of myosin light-chain kinase with ML-7 blocked this RhoA-induced increase in extrusion rate (Figure 2C), indicating that it depends on myosin contractility. Importantly, division rate was not higher in Opto-RhoA than Opto-Ctrl cells, either illuminated or not (Figure 2D). Accordingly, mitosis-blocking Mitomycin C treatment did not reduce the RhoA-dependent induction of extrusions (Figure 2C), showing that compensation of a potential effect of RhoA on proliferation^43^ is not responsible for the observed effect. We found that in Opto-RhoA illuminated regions, tensile stress increased within a approximately 1–2 cell diameters-region surrounding the extruding cell, 60 to 30 minutes prior to its extrusion (Figure 2 E-G). Following this peak, tensile stress decreased rapidly, coinciding with the elimination of the cell and the convergent motion of neighboring cells to close the gap. In contrast, Opto-Ctrl cells exhibited a different mechanical profile: extrusions were preceded by a sustained increase in compressive stress (negative stress values) lasting more than two hours (Figure 2 E,F,H), in line with previous reports^2,9,10^. This suggests that the extrusion of highly contractile cells arises from mechanical tension within the monolayer. Supporting this view, optogenetic Rac1 activation, which reduced local tension, decreased the extrusion rate (Figure S6). Interestingly, whereas most extruded cells in Opto-Ctrl or non-illuminated Opto-RhoA regions were apoptotic, the majority of tension-induced extrusions showed no active caspase-3 staining (Figure 2 I,J). These findings reveal that tension can substitute for compression in driving extrusion, raising the possibility of a mechanistically distinct pathway for eliminating highly contractile cells.

**Figure 2:**
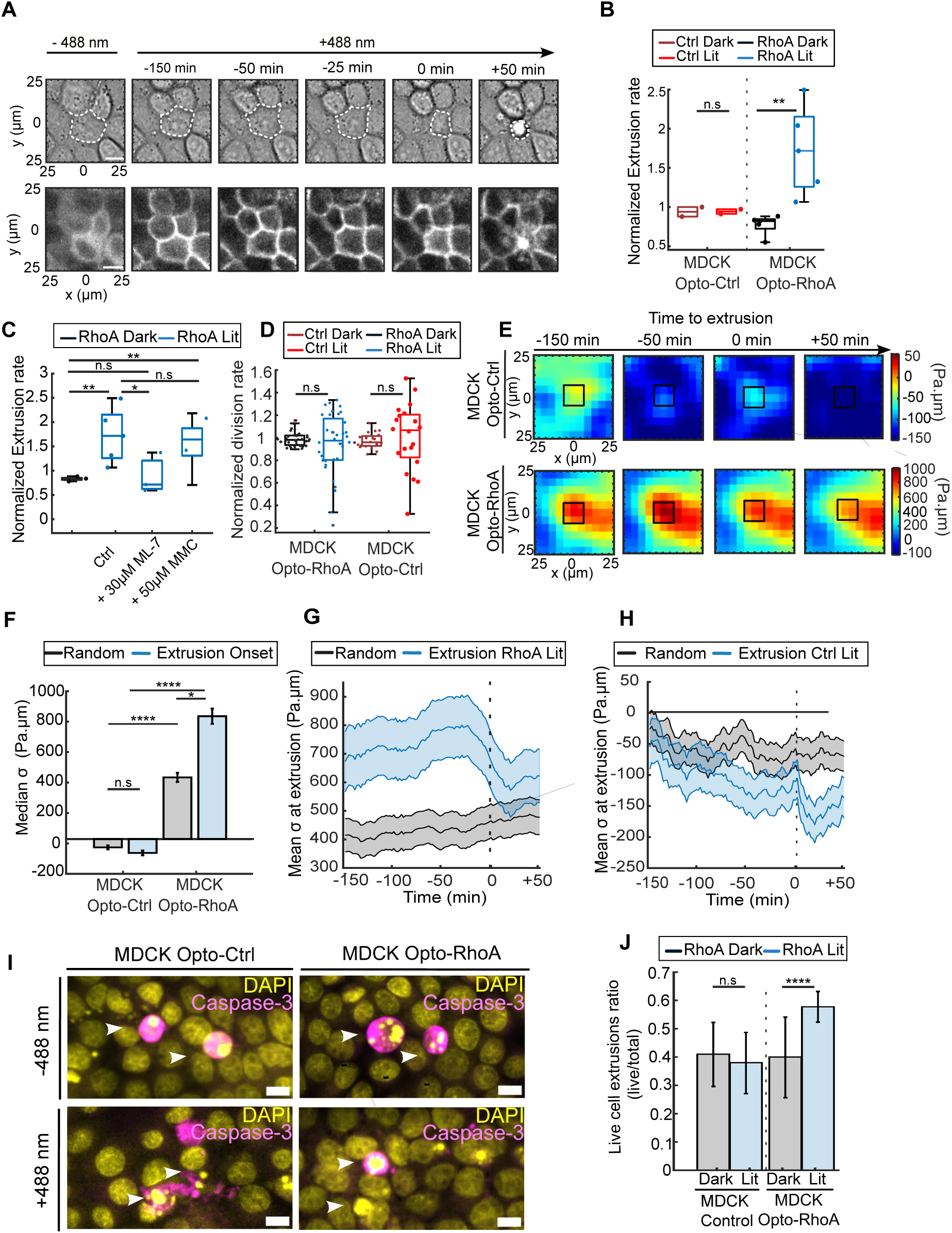
RhoA-induced tensile stress generates cell extrusions. **A** Cell extrusion event out of an MDCK Opto-RhoA monolayer before (−488 nm) or after the start of stimulation (+488 nm). (Top): brightfield, (bottom): CRY2-mCherry signal. Cell extrusion onset is defined as the moment cell area starts shrinking (0 min). **B** Extrusion rate in the stimulated (Lit) or non-stimulated (dark) regions normalized by their area sizes for Opto-Ctrl (N=2) and Opto-RhoA cells (N=5). Wilcoxon rank-sum test: n.s. p = 0.9681; ** p = 0.0079 from left to right. **C** Normalized extrusion rate for untreated cells (Opto-RhoA control, N=5 n=51) and cells treated with 30µM of ML-7 to block contractility (N=3 n=31) or 50 µM MMC to block proliferation (N=2 n=30). Wilcoxon rank-sum test: ** p = 0.0079; n.s., p = 0.1429; ** p = 0.0043; * p = 0.0357; * p = 0.0380; n.s., p = 0.8355, from left to right. **D** Division rate in Opto-RhoA (N=2 n=35) and Opto-Ctrl (N=2, n=22) cells in the stimulated (Lit) or unstimulated area (Dark), normalized to the total number of cell divisions in each condition. Wilcoxon rank-sum test: n.s., p = 0.8969; n.s., p = 0.4407, from left to right. **E** Heatmaps of the evolution of mean isotropic stress of Opto-Ctrl (top, N=2 n=94 cells) and Opto-RhoA cells (bottom, N=3, n=116 cells) within a 50 × 50 µm window centered around the extrusion locations (dark square). **F** Median isotropic stress in region of 10 x 10 µm centered at extrusion sites in stimulated areas (blue) or random locations (dark). Wilcoxon rank-sum test: n.s., p = 0.1643; **** p = 1 × 10⁻⁸; **** p = 6.27 × 10⁻¹⁰; * p = 0.0390 from left to right. **G, H** Mean isotropic stress at extrusion sites (blue) within stimulated areas or random locations (black) for Opto-RhoA cells (G) or Opto-Ctrl (H) respectively. Time = 0 represents the onset of extrusions. **I** Representative images of cell extrusions (white arrow-head) of Opto-RhoA or Opto-Ctrl cells light-stimulated (+488 nm) or unstimulated (−488 nm) and stained for active Caspase-3 (magenta). **J** Ratio of caspase-negative cell extrusions in MDCK Opto-RhoA and Opto-Ctrl cell lines, unstimulated (gray bars) or stimulated for 8 hours (blue). (N=2 n=408 cells, N=2 n=242 cells, N=2 n=385, N=2 n= 276 cells respectively). Wilcoxon rank-sum test : n.s., p =0.5129; **** p = 2.61 × 10⁻⁷ from left to right. Box plots represent the median, interquartile (box), and 1.5 IQR (whiskers). Curves represent the mean +/- s.e.m. See also Figure S3, S4, S5, S6 and Video S2.

To question how tension could trigger cell extrusion, we turned to our 3D phase-field model. As before, we simulated the mechanical effects of RhoA activation by increasing cellular stiffness within the activated region. We varied the stiffness contrast between activated and non-activated regions in our simulations, and found that the probability of cell extrusion increased with the effective stiffness of the activated cells (Figure 3A). Local averaging of in-plane isotropic stress around extrusion events revealed a clear dichotomy: extrusions arising in stiffened regions correlated with elevated in-plane tension, whereas extrusions in non-activated regions were preceded by compression (Figure 3 B,C). This matched our experimental observations. However, the physical link between in-plane stress and extrusion is not immediately obvious. To better understand this mechanism, we leveraged the 3D nature of our model to examine the out-of-plane stress component, σ_zz_, since extruding cells are displaced vertically from the monolayer. Firstly, we found that the magnitude of σ_zz_ was significantly higher in activated cells that subsequently extruded than in activated cells that did not extrude (Figure S7B), demonstrating that increased out-of-plane loading is associated with extrusion. However, this increase may simply be a mechanical readout of the extrusion process and is not directly related to the enhanced in-plane tension. We therefore asked which mechanical features arising from the enhanced in-plane tension are linked to cell extrusion. A key observation of the simulations is that temporal fluctuations in σ_zz_ (quantified via the coefficient of variation over time at each spatial location) were enhanced within the activated region (Figure 3D).

**Figure 3:**
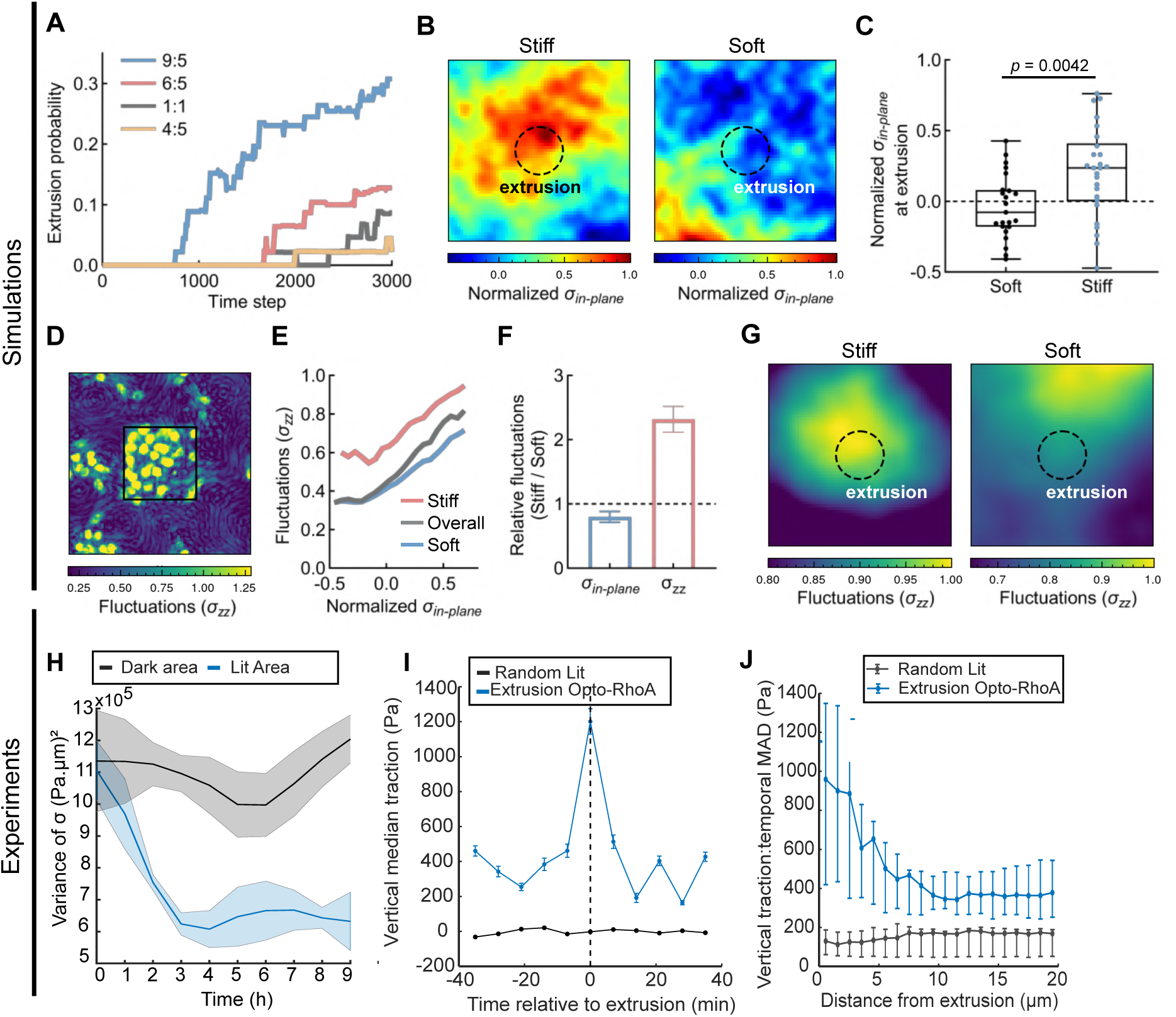
Stiffness-dependent amplification of out-of-plane stress fluctuations promotes tension-induced cell extrusion. (**A-G** Results from simulations.) **A** Simulated extrusion probability increases with the stiffness ratio between activated and non-activated cells. Each curve represents the mean across 3 independent simulations. **B** Averaged maps of normalized in-plane isotropic stress (*σin-plane*) around extrusion events. The dashed circle indicates the extrusion region, defined by the mean cell radius. **C** Quantification of mean *σin-plane* within the extrusion region confirms that extrusions in the stiff (activated) domain occur under significantly higher tensile stress than those in soft regions (p=0.0042). Box plots represent the median (center line), interquartile range (box), and minimum to maximum values (whiskers). **D** Spatial map of vertical stress fluctuations (*σzz*), quantified as the temporal coefficient of variation, shown for the same simulation and time window as in Figure S2C. **E** *σzz* fluctuations plotted against local *σin-plane* (3 independent simulations). **F** Relative amplitude of *σin-plane* and *σzz* fluctuations in stiff versus soft regions (3 independent simulations, mean ± SD). **G** Averaged spatial maps of *σzz* fluctuations centered on extrusion sites demonstrate localized amplification of vertical stress fluctuations in stiff regions (left), but not in soft regions (right). (Quantifications in Figure S7A-C). (**H** Results from 2D TFM experiments) H Mean variance of *σin-plane* within the stimulated (blue, 150 × 150 µm illumination region) and non-stimulated (dark) areas (N=2). (**I-J** Results from 2.5D TFM experiments) **I** Median vertical tractions (*tz*) computed at extrusion sites over time, using a 5 x 5µm window centered on extrusion events (blue, N=2 n=38, maps in Figure S7D) or at random positions (black N=2, n=45). **J** Temporal fluctuations of vertical traction forces around the extrusion events (blue) or at random positions (grey). Error bars are 95 % bootstrap confidence intervals of that median (1000 resamples). See also Figure S7.

To probe the origin of these fluctuations, we computed the correlation between in-plane isotropic stress and σ_zz_ fluctuations across all spatial points in the monolayer. We found that sites of high in-plane tension exhibited stronger σ_zz_ fluctuations across the monolayer (Figure 3E, gray curve), indicating that accumulated tension can amplify vertical stress variability. Given the mechanical heterogeneity of the tissue, we asked whether this amplification differs between stiff (activated) and soft (non-activated) cells. When separating the monolayer into these regions, the correlation was significantly stronger in the stiff domain. This suggests that stiffer cells, which resist in-plane deformation, more effectively redirect the built-up tension into elevated out-of-plane stress fluctuations (Figure 3F, red bar).

Since stiff cells naturally accumulate in-plane tension (as shown in Figure S2) and more efficiently transfer it into vertical fluctuations, we hypothesized that these enhanced σ_zz_ fluctuations act as direct mechanical precursors to extrusion in the activated region by signaling local out-of-plane instability. To test this, we averaged the simulated local out-of-plane stress fluctuations found around extrusion events. Remarkably, in the activated region, extrusion locations significantly overlapped with areas of elevated σ_zz_ fluctuations, whereas this association was much weaker and more spatially diffuse in the non-activated region (Figure 3G, Figure S7C). This suggests that enhanced out-of-plane stress fluctuations may serve as a mechanical cue for extrusion in stiff cells. In contrast, activated regions showed suppressed in-plane isotropic stress fluctuations (Figure 3F, blue bar), consistent with reduced deformability of stiff cells. To experimentally validate the mechanical mechanism predicted by the model, we quantified stress fluctuations in stimulated and non-stimulated regions of the monolayer. Consistent with the simulation results, activation of contractility and the associated tissue stiffening strongly reduced fluctuations in in-plane isotropic stress (Figure 3H).

To further decipher the mechanisms underlying tension-induced extrusion, we next sought to experimentally probe the out-of-plane mechanical component predicted by the model. While the simulations quantify internal vertical stress fluctuations (σ_zz_), these quantities are not directly accessible experimentally. We therefore used 2.5D traction force microscopy^44,45^ as a proxy to measure vertical traction forces and their temporal fluctuations during extrusion events in globally stimulated confluent monolayers of MDCK Opto-RhoA cells. Consistent with the model predictions, whereas in-plane isotropic stress fluctuations were reduced upon RhoA activation, vertical forces emerged prior to extrusion events, with vertical traction force fluctuations significantly enhanced at extrusion sites (Figure 3I, S7D). Importantly, this mechanical signature was specific to tension-induced extrusion events, as the same analysis performed at randomly selected positions within the stimulated monolayer did not reveal any comparable spatial or temporal pattern (Figure 3J). Together, these findings experimentally support a model in which RhoA-associated increased cellular stiffness promotes extrusion by redirecting accumulated in-plane tension into out-of-plane mechanical fluctuations

### Cell extrusion is driven by tensile stress amplitude rather than mechanical mismatch

Previous studies have shown that local mechanical mismatches can drive cell extrusions^46,47^ in a manner reminiscent of cell competition^18,48,49^. In contrast, our model predicts that high tension amplitude alone is sufficient to trigger extrusion, without requiring mismatch (Figure 3 A-C). Accordingly, non-illuminated Opto-RhoA cells show a higher extrusion rate than Opto-Ctrl cells that are less contractile (Figure S5), suggesting that absolute tension level, without local mismatches, correlates with cell elimination. To investigate further whether cell extrusion is predominantly influenced by the magnitude of tensile stress or by local stress gradients, we modulated the geometry of the stimulated regions. We first varied the size of the stimulated areas (Figure 4A, Video S3), reasoning that smaller square patterns, due to their higher perimeter-to-area ratios, would generate stronger stress gradients at the interface between stimulated and non-stimulated regions. In contrast, larger patterns were expected to display a more uniform stress distribution. Interestingly, Opto-RhoA cells stimulated with smaller illumination patterns (44 x 44 µm and chessboard patterns of five 50 x 50 µm squares) showed a higher extrusion probability than cells stimulated within larger regions (150 x 150 µm and 260 x 260 µm) (Figure 4B, Figure S8 A-E). Importantly, these smaller regions also exhibited higher average isotropic stress (Figure 4C), making it unclear whether gradients or absolute tension were the key driver.

**Figure 4:**
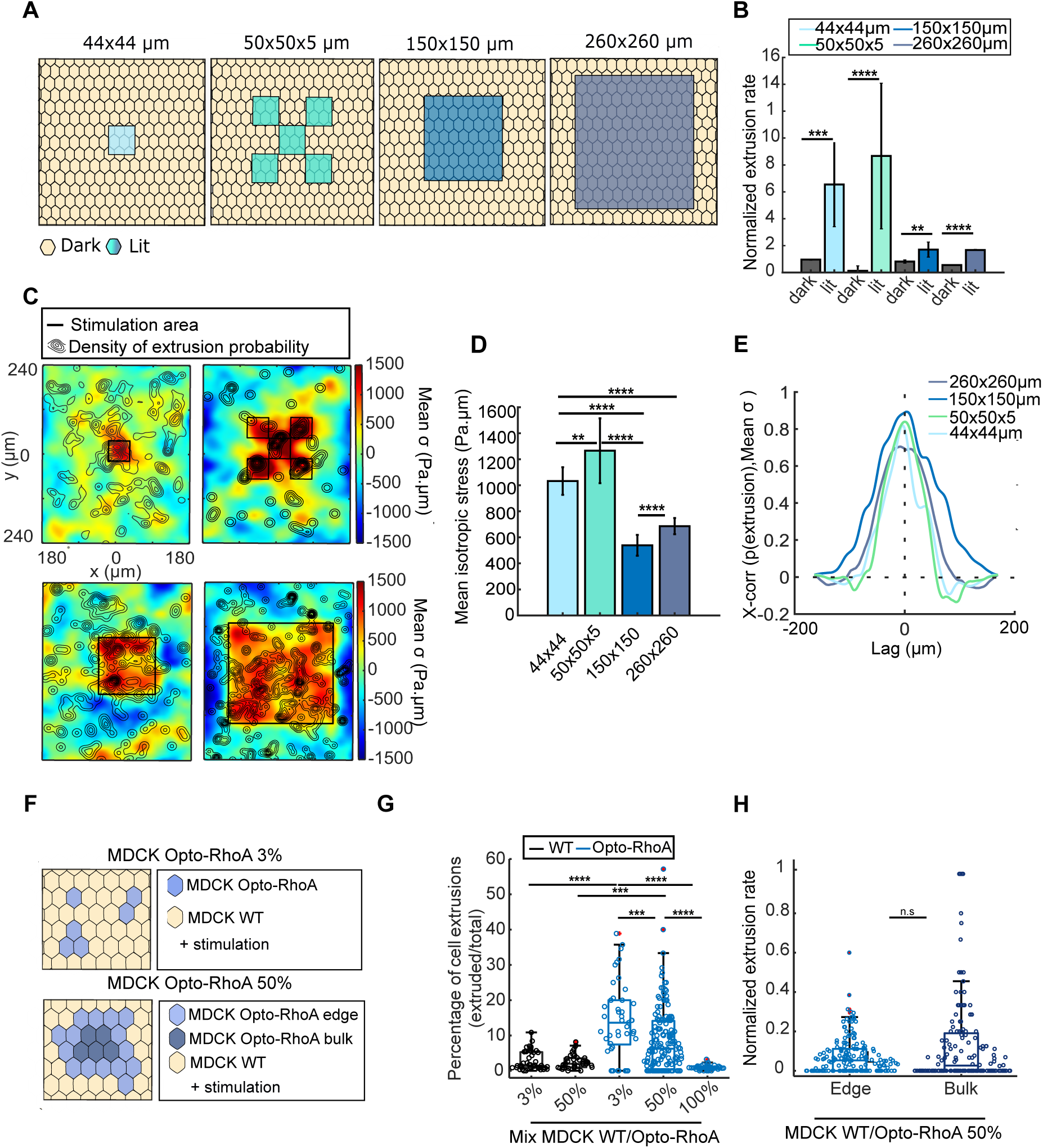
RhoA-mediated extrusion is primarily driven by tensile stress amplitude. **A** Modulations of the size and shape of the illuminated region. **B** Normalized extrusion rate of Opto-RhoA cells within (lit) or outside (dark) the illuminated regions (N=3 n=339, N=2 n=132, N=5 n=115, N=2 n=259 events from left to right). Wilcoxon rank-sum test: *** p = 7.10 × 10⁻⁴; **** p = 1.82 × 10⁻⁶; ** p = 0.0079; **** p = 8.61 × 10⁻⁸ from left to right. **C** Density maps of extrusion locations (black level curves) and mean isotropic stress (color coded) in the increasing order of regions stimulation size (from left to right, black squares). Related to Figure S8F-I. **D** Average *σin-plane* inside the stimulated areas (lit): 44 x 44 µm (light blue, N=3), 50 x 50 µm x 5 (cyan, N=2), 150 x 150µm (blue, N=2) and 260 x 260 µm (dark blue, N=2). Wilcoxon rank-sum test: **** p = 1.34 × 10⁻¹⁵; **** p = 1.05 × 10⁻²⁰; ** p = 0.001521; **** p = 1.89 × 10⁻¹⁵; **** p = 6.54 × 10⁻⁷ from top to bottom and left to right. **E** Normalized cross-correlation functions between the spatial distribution of cell extrusion probability and isotropic stress in these different conditions. **F** Representative scheme of co-culture experiment. MDCK WT cells (yellow) were mixed with 3% (top) or 50% (bottom) MDCK Opto-RhoA (blue) cells. Light blue: Opto-RhoA cells localized at the interface with WT cells. Dark blue: with no WT neighbors. **G** Percentage of extruded cells in co-culture experiments with 3%, 50% Opto-RhoA/WT cell proportions or pure culture of Opto-RhoA cells (N=3). Wilcoxon rank-sum test: **** p = 9.89 × 10⁻¹¹; **** p = 3.01 × 10⁻⁷; *** p = 5.43 × 10⁻⁴; *** p = 4.40 × 10⁻⁴; **** p = 1.38 × 10⁻⁶, from top left to bottom right. **H** Normalized extrusion rates at the edge or within the bulk of Opto-RhoA cell clusters in 50% Opto-RhoA/WT co-cultures (N=3). Related to Figure S9. Wilcoxon rank-sum test: n.s., p = 0.6164. Bar graphs represent the mean +/- s.e.m. Box plots represent the median, interquartile (box), and 1.5 IQR (whiskers). See also Figure S8, S9 and Video S3.

To distinguish between these possibilities, and to assess whether extrusion localization correlates more closely with stress magnitude or with spatial gradients, we next mapped the spatial distribution of isotropic stress together with extrusion probability. Strikingly, extrusion events occurred preferentially within regions exhibiting the highest tensile stress inside the stimulated domain (Figure 4D). This observation was further confirmed by cross-correlation analysis between local extrusion probability and average isotropic stress values, which revealed a strong, center-peaked correlation across all stimulation sizes (Figure 4E, Figure S8 F-M), highlighting that extrusion probability robustly correlates with stress amplitude. In contrast, cross-correlation analysis between extrusion probability and stress gradients generated a weaker and broader correlation peak (Figure S8 J-M), revealing a modest spatial offset between regions of high stress gradient and extrusion sites (Figure S8 N,O).

Furthermore, cells undergoing extrusion did not display elevated levels of ArhGEF11-CRY2-mCherry relative to neighboring cells prior to optogenetic stimulation (Figure S8 P,Q). Quantification of ArhGEF11-CRY2 fluorescence intensity at extrusion sites revealed only an increase of signal coincident with cell area reduction prior to rounding up and detachment, thus when cells were already engaged in extrusion (Figure S8 Q,R). These results indicate that extrusion events are not associated with locally elevated expression of the optogenetic construct and therefore cannot be explained by cell-to-cell variability in ArhGEF11-CRY2-mCherry expression. Together, these analyses demonstrate that extrusion probability is better predicted by stress amplitude than by stress gradients.

To test the contribution of local mechanical mismatches, we mixed MDCK Opto-RhoA and WT MDCK cells to generate stable and cell-type specific stress mismatches. When co-cultures contained 3% Opto-RhoA cells this population formed small clusters (1-5 cells) all sharing adhesive junctions with WT cells. In co-cultures containing 50% Opto-RhoA cells, they formed larger clusters (36 cells on average), with some cells not sharing any junctions with their WT counterparts (bulk, Figure 4F). Opto-RhoA cells within smaller clusters displayed a significantly higher probability to extrude than cells in larger clusters or in pure Opto-RhoA cultures (Figure 4G), consistent with our previous results showing higher extrusion probability and tensile stress in smaller stimulated regions (Figure 4D). In addition, within mixed clusters, cells close to the interface were not significantly more prone to extrusion, regardless of whether they were WT or Opto-RhoA cells (Figure 4H, Figure S9). Consistently, WT cells did not exhibit increased extrusion rates in co-culture conditions. These results collectively indicate that while local mechanical mismatches may play a permissive role by modulating the local stress as suggested by our numerical model (Figure S2 G,H), it is primarily the magnitude of tensile stress that determines the likelihood of extrusion.

### Cytoskeletal remodeling under tension triggers cell extrusion

Altogether, our modeling and experimental results indicate that local cell stiffening mediated by contractility leads to the accumulation of tensile stresses. Those are redirected into amplified out-of-plane mechanical fluctuations, which are associated with extrusion. Given the central role of RhoA in stress fiber assembly and actomyosin contractility, we reasoned that this mechanical behavior likely depends on actomyosin structures capable of sustaining and transmitting forces across cells. We thus investigated whether tension-induced extrusion is associated with a specific organization of the actomyosin cytoskeleton. Following uniform RhoA activation across the MDCK Opto-RhoA monolayer, extruding cells displayed a strikingly distinct cytoskeletal organization compared to neighboring non-extruding cells, with a strong enrichment of actin fibers decorated with phospho-myosin (Figure 5 A,B). Although we observed a noticeable reinforcement of the apical actomyosin belt, the most pronounced enrichment occurred at the basal side, within stress fibers and along the cell periphery (Figure 5 C,D). Interestingly, apical phospho-myosin levels were lower in extruding cells than in their neighboring cells, suggesting that surrounding cells participate in the formation of a transcellular myosin ring. Live-cell microscopy further revealed that F-actin accumulated apically throughout the extrusion process both in the extruding cell and its direct neighbors (Figure 5E), suggesting a purse-string-like mechanism driving the expulsion of the center cell^7,19,20^. In contrast, F-actin progressively accumulated basally prior to extrusion before abruptly decreasing approximately 50 minutes before cell detachment (Figure 5E). Stress fibers appeared markedly more aligned in cells prior to their extrusion (Figure 5 F-H, Video S4 A-B). Active gel theory predicts that actomyosin contractile stress scales with the nematic order parameter of the cytoskeleton^50^, such that increased actin alignment amplifies anisotropic contractility. Consistent with this idea, actin fiber alignment has been implicated in large-scale mechanosensing and in the buildup of substantial traction forces at the cell–ECM interface^51^. Thus, the progressive basal actin alignment observed before extrusion is expected to locally increase actomyosin-dependent tensile stress within the extruding cell. Accordingly, actin alignment increased progressively (Figure 5H), in parallel with the rise in tensile stress that peaks approximately 50 minutes before cell detachment (Figure 2G).

**Figure 5:**
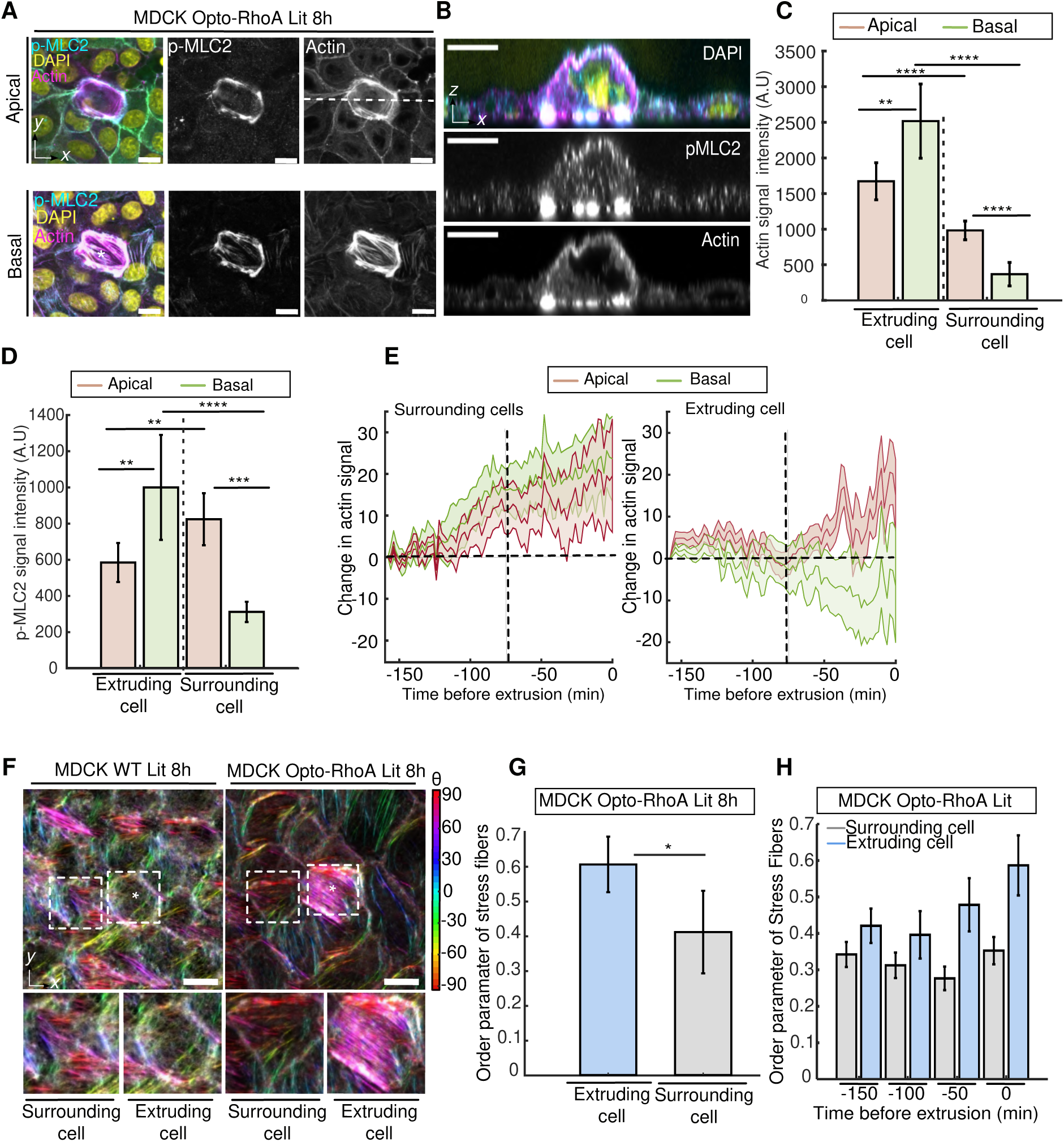
Specific organization of cytoskeleton in tension-induced cell extrusion. **A** Apical (top) and basal (bottom) views of a cell undergoing extrusion (*) within stimulated Opto-RhoA cells. Magenta: actin, Cyan: p-MLC2, yellow: nuclei. **B** Transversal view from **(A)**. **C,D** Quantification of mean actin and p-MLC2 fluorescence intensities at the apical (red) and basal (green) planes in MDCK Opto-RhoA extruding (N=2, n=15) and surrounding cells (N=2, n=84). Wilcoxon rank-sum test. **C**: ** p = 0.001404; **** p = 1.61 × 10⁻⁶; **** p = 8.58 × 10⁻¹⁰; **** p = 1.82 × 10⁻¹³, from left to right. **D**: * p = 0.01440; ** p = 0.001945; **** p = 1.40 × 10⁻⁷; **** p = 7.84 × 10⁻¹⁷, from left to right. **E** Quantification of the temporal changes in basal (green) and apical (red) actin signal in extruding cells (right, N = 3 n=11 cells) and their neighboring cells (left, N=3 n=11). **F** Orientation of actin stress fibers at the basal side of MDCK WT (left) and MDCK Opto-RhoA (right) monolayers illuminated for 8 hours with blue light (488nm). *: extruding cell. Bottom: details of the surrounding (left) or extruding cell (right). **G** Mean order parameter of stress fibers orientation in MDCK Opto-RhoA extruding (blue; N = 2, n = 15) and surrounding cells (gray; N = 2, n = 15). Wilcoxon rank-sum test: * p = 0.03814. **H** Temporal evolution of the actin stress fibers order parameter in MDCK Opto-RhoA extruding (blue) and surrounding cells (gray). Scale bars: 10µm. Bar graphs: mean +/- s.e.m. See also Figure S10 and Video S4.

To further investigate the functionality of stress fibers in tension-induced extrusion, MDCK Opto-RhoA cells were cultured on soft PDMS substrates (2 kPa), a condition known to reduce stress fiber formation. Basal F-actin signal intensity in stimulated cells was significantly lower on soft substrates than on stiffer 15 kPa PDMS substrates (Figure S10 A-B), with a reduced stress fiber density and a marked decrease in RhoA-induced extrusion probability (Figure S10C-D). In addition, we compared cytoskeletal organization in small activated regions (44 × 44 µm), where tissue stress and extrusion probability are high, with larger activated regions (260 × 260 µm), where both tissue stress and extrusion probability are reduced (Figure S10 E-F). Small activated regions displayed a strong increase in basal actin intensity following stimulation, whereas larger regions exhibited a weaker response after stimulation with blue light (Figure S10 E-H). These observations suggest that local confinement of contractility promotes more robust actin remodeling and stress fiber assembly, which results in higher tissue tension, cell stiffening and extrusion proficiency. Together, these findings show that tensile stress buildup drives basal actin cytoskeleton accumulation and remodeling that culminates in cell detachment and extrusion.

### RhoA-mediated contractility induces live basal cell extrusion

The emergence of prominent basal stress fibers prompted us to investigate whether increased cell-matrix coupling could favor basal extrusion, as previous studies have shown that strengthening of the cell-matrix interface can facilitate extrusion towards the basal side^11,14,52,53^. We thus cultured MDCK Opto-RhoA monolayers on fibrillar collagen I matrices. Under these conditions, cells could extrude both apically and basally (Figure 6 A,B). While Opto-Ctrl cells were predominantly extruded towards the apical side, Opto-RhoA cells displayed a higher proportion of basal extrusions that was enhanced under illumination, up to 60% (Figure 6 B,C). To rule out that the basal bias of extrusion resulted from our optogenetic strategy, particularly from non-physiological apical or basal localization of contractility, we stimulated WT cells with CN03, which also enriched basal extrusions (Figure S11C). Two additional optogenetic tools were further used to control the activation of RhoA along the apico-basal axis. We expressed ArhGEF11-CRY2-mCherry with either ITB1-CIBN-GFP or N-shroom-CIBN-GFP allowing the recruitment of ArhGEF11, and thus RhoA activation, to the basal and apical sides, respectively^54^ (Figure S12 A,B). After illumination, cells expressing either one or the other both showed increased extrusion rates and basal extrusion proportions (Figure S12 B,C), suggesting that it is the overall increase in contractility — rather than the location of the contractile mechanism — that causes basal extrusions. We observed that both optogenetic and CN03-mediated activation of RhoA triggered predominantly non-apoptotic extrusions on glass (Figure 2 I,J, Figure S11D). However, since basal extrusions were not possible in these conditions, we asked whether extrusion directionality (apical *versus* basal) was linked to the fate of extruded cells (dead *versus* alive) in permissive conditions. To investigate this, we cultured Opto-Ctrl and Opto-RhoA cells on collagen and monitored apoptosis during live-cell imaging (Figure 6D). Annexin V signal was detected shortly after the onset of extrusion for apical extrusion events in both Opto-Ctrl and Opto-RhoA cultures, with an average delay of 20 and 2 minutes, respectively (Figure 6E, Video S5). In contrast, Annexin V signal appeared substantially later for basal extrusion events, with an average delay of 40 minutes (Figure 6E). Thus, the initiation of basal extrusion is not directly associated with apoptotic commitment. Nevertheless, most basally extruded cells eventually die within the matrix within hours after extrusion. One possible explanation is that collagen-1 alone does not provide sufficient survival signals to sustain viability of basally extruded cells.

**Figure 6:**
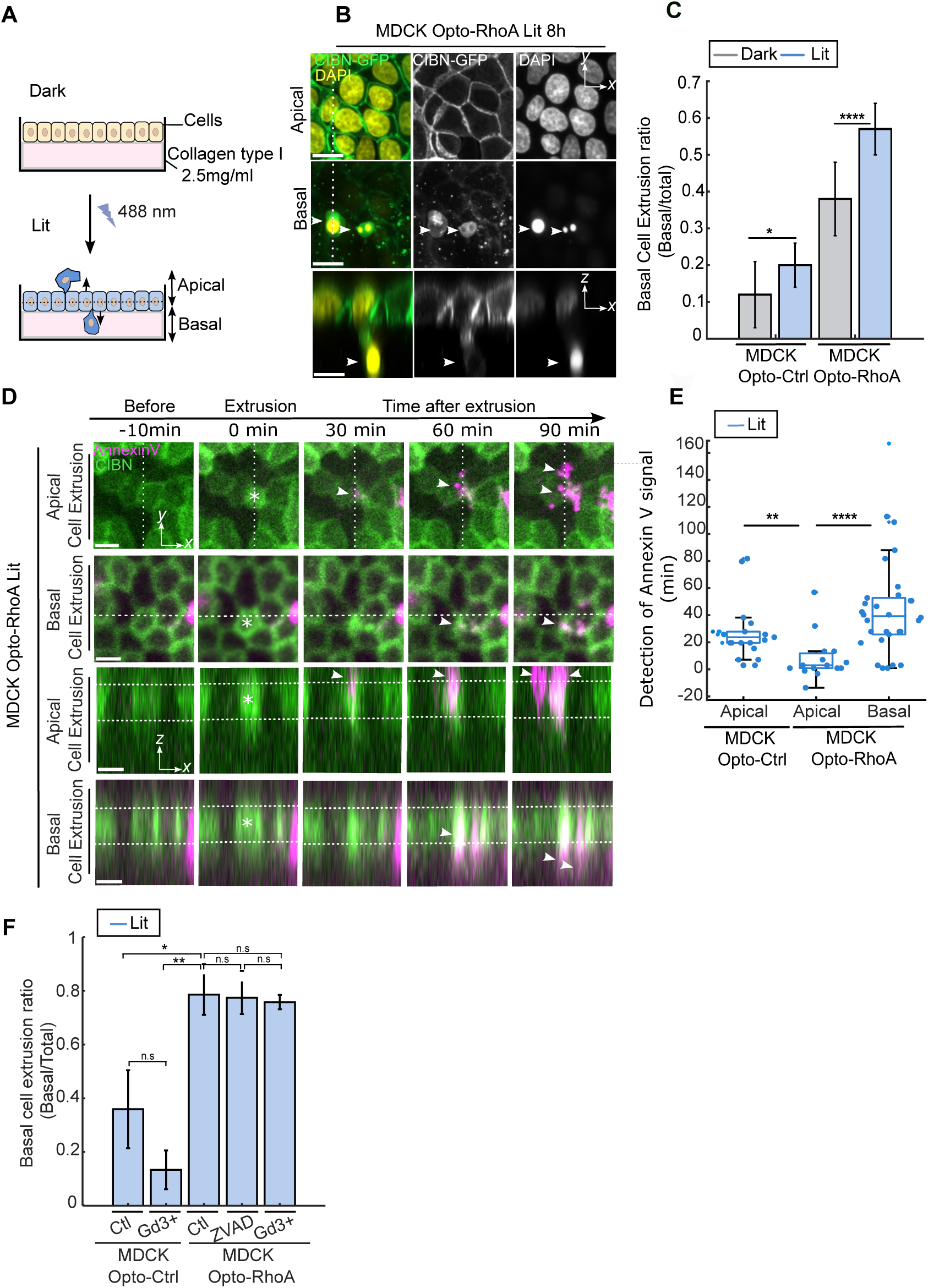
RhoA-mediated contractility induces live basal cell extrusion. **A** MDCK monolayers cultured on a collagen type I matrix (2.5 mg/mL) enabling both apical and basal extrusions. **B** Basal extrusion of Opto-RhoA cells (arrowheads) cultured on a collagen type I matrix following 8 h of optogenetic stimulation, at the apical junction plane (top) and basal plane (middle). Bottom: transversal XZ view. **C** Ratios of basal cell extrusion (basal/total cell extrusions) in MDCK Opto-Ctrl and MDCK Opto-RhoA cells, without stimulation (dark, gray bars) (N=2 n=194 cells, N=2 n=139) or after 8 hours of stimulation (Lit 8h, blue bars) (N=2 n=168, N=2 n=334 cells respectively). Wilcoxon rank-sum test: * p = 0.01705; *** p = 9.09 × 10⁻⁴, from left to right. **D** Detection of apoptosis by live-cell imaging of Annexin V fluorescence (magenta) in apically (top) or basally (bottom) extruded MDCK Opto-RhoA cells. White arrows indicate the onset of Annexin V fluorescence detection. Time = 0 corresponds to the beginning of the cell extrusion event. Dashed lines delineate the height of the epithelial monolayer. **E** Median temporal offset between extrusion initiation and Annexin V signal detection for apical and basal extrusions of illuminated MDCK Opto-Ctrl (N = 1, n = 21) and Opto-RhoA cells (N = 3, n = 105). Wilcoxon rank-sum test: *** p = 4.56 × 10⁻⁴; *** p = 1.44 × 10⁻⁴, from left to right. **F** Basal cell extrusion ratios for MDCK Opto-Ctrl and MDCK Opto-RhoA cell lines without treatment (Ctl) or treated with 50 µM Gadolinium chloride (Gd) or ZVAD-FMK (N=2 n=35, N=1 n=16, N=3 n=90, N=2 n=61, N=2 n=60 cells respectively). Wilcoxon rank-sum test: * p = 0.0423; n.s., p = 0.5357; ** p = 0.01482; n.s., p = 0.4054; n.s., p = 0.1587, from top to bottom and left to right. Bar graphs represent the mean +/- s.e.m. Box plots represent the median, interquartile (box), and 1.5 IQR (whiskers). See also Figure S11, S12, S13 and Video S5.

Compression-induced apical extrusion, both apoptotic and alive, has been shown to depend on the Sphingosine-1-Phosphate (S1P) pathway downstream of the stretch-activated ion channel Piezo-1^55,56^. We thus questioned whether activation of this pathway is also required for the live basal extrusions induced by RhoA-mediated tension. Interestingly, neither the Piezo-1 inhibitor Gd³⁺, nor caspase inhibition using Z-VAD-FMK treatment significantly reduced basal extrusion rates or affected the proportion of basal cell extrusions in stimulated MDCK Opto-RhoA monolayers (Figure 6F, Figure S13). If anything, the inhibition increased the total extrusion rate, which could be due to increased contractility, as recently suggested^57^. Together, these findings suggest that RhoA-mediated tensile stress induces a distinct mode of basal extrusion that is largely independent of Piezo-1 signaling and apoptosis.

## Discussion

Several studies have previously shown that compressive stress triggers cell extrusion *in vivo* and *ex vivo*^2,9,10,19,58,59^. Here, we reveal an unexpected, alternative pathway: high RhoA activity generates intercellular tensile stress that is sufficient to trigger extrusion. In RhoA-activated regions, elevated tension promotes out-of-plane stress fluctuations, amplified by increased stiffness and actin cytoskeletal remodeling, which are associated with extrusion. Elimination of apoptotic cells in MDCK monolayers, Drosophila larval epithelium and zebrafish larval epidermis is known to depend on local actomyosin accumulation at cell-cell junctions downstream of the RhoA/ROCK pathway during its final stage^6,16,17,53^. We show that RhoA activation and elevated contractility act upstream to initiate extrusion. This is consistent with previous observations that high RhoA-dependent contractile activity enhances extrusion rates in MDCK cells^60,61^ and in the zebrafish epidermis^56^. Similarly, larval epidermal cells in the abdominal epidermis of Drosophila melanogaster preferentially extrude in regions of high cortical tension^62^. While these reports pinpointed the role of myosin as a potential precursor of extrusion, they did not provide a physical description of the mechanical landscape associated with these events, nor did they provide a model explaining the mechanistic role of contractile forces. Importantly, a recent article showed that apical cell extrusion at the tip of intestinal villi results from contractile tension^63^, challenging the classical view that cell shedding results from tissue crowding in this epithelium. Our results confirm the role of mechanical tension as an initiator of cell extrusion and provide an unprecedented precision in mechanical force measurement, allowing us to propose a physical explanation for this phenomenon. On the contrary, it has been shown that intercellular tension opposes apoptotic cell extrusion in CaCo2 cells^64^ and that relaxation of cell contractility in the vicinity of apoptotic cells is required for their extrusion^8^. This ambivalence might be due to the difficulty of precisely actuating and measuring mechanical forces simultaneously. Optogenetic modulation of RhoA activation allowed for quantitative, spatially controlled modification of the mechanical landscape of the tissue, leading to a local or global increase in intercellular tension, resulting both in increased extrusion rates.

Another possible explanation for these different findings is that the physiological context (cell differentiation, ECM properties, etc.) and overall mechanical environment (beyond intercellular mechanical stress) might modulate the cellular response to tension *vs.* compression. Indeed, we observed that compressive stress is responsible for extrusions in normal cells, while intercellular tension drives extrusions in stiffer ones. It is worth noting that the tensile stress values we measured (500-1000 Pa.µm after RhoA activation) are comparable to what has been measured in similar epithelial models^65,66^ and lower than in patient-derived tumors^18^. This suggests that both the cell-autonomous contractility and the local mechanical landscape at the multicellular level determine the cellular ability to cope with tension.

Our numerical model provides a minimal framework to connect these experimental observations. In the simulations, the RhoA-induced change in cell mechanics is represented by increasing γ, which modulates interfacial mechanics and thereby the effective deformability of the cells^41,42^. This representation is consistent with our AFM measurements showing increased cell stiffness following RhoA activation, and reproduces the experimentally observed buildup of in-plane tension, reduced cell motion, and enhanced out-of-plane mechanics associated with extrusion. Regarding the latter, both the magnitude and the fluctuations of vertical stress are elevated in extruding cells, and our data cannot fully disentangle their contributions. The increased magnitude may, however, simply be a mechanical readout of the extrusion process itself, since a cell leaving the monolayer necessarily experiences a net vertical load. We also note, however, that active cortical contractility is not implemented as a separate contribution in the present model. Accordingly, the γ-dependent results should be interpreted as a coarse-grained representation of the RhoA-induced change in effective interfacial mechanics, rather than as evidence that stiffness acts independently of cortical contractility or as a literal description of cortical tension at every subcellular location.

Remarkably, our results demonstrate that elimination of RhoA cells happens more frequently when less-contractile cells surround hypercontractile ones, reminiscent of a non-autonomous process described in previous studies^8,18^. Although in intestinal villi extrusion typically involves the less contractile cells in direct contact with more contractile neighbors^63^, we found that it is the RhoA-activated MDCK cells that exhibited the higher probability of extrusion when mixed with WT cells. This effect occurred regardless of their distance from the interface, suggesting that the involved mechanism is different from non-autonomous, cell-competition situations observed in intestinal villi^63^ or between MDCK cells expressing different E-cadherin levels^18^. Instead, cell-autonomous RhoA-induced stiffness, which we found to be higher when small subsets of cells show elevated contractility, is sufficient to explain cell extrusion as predicted by our numerical model. In addition, elevated tension at the whole-population level was accompanied by higher extrusion rates. Altogether, while we cannot fully exclude that local tension mismatches also participate in this process, our results show that absolute tension levels are sufficient to drive extrusion.

At the molecular level, we observed that RhoA activation led to an enrichment of apical and basal actomyosin networks, accompanied by reduced E-cadherin at cell-cell junctions. Our numerical model predicts that the stiffness increase due to actin accumulation or re-arrangement is sufficient to induce extrusions. However, while E-cadherin reduction is not necessary in this scenario, it could participate in stress dissipation. Our analysis of stress fiber dynamics shows that the actin cytoskeleton responds to increased tension and rigidity through local stress fiber alignment before extrusion. We propose that this increase in nematic actin order amplifies contractile stress at the basal surface, consistent with previously shown links between large-scale actin-based mechanosensing^51,67^, stress fiber alignment, increased traction stress and cell stiffening^68,69^, until the transmitted load exceeds the capacity of cell–ECM adhesions to sustain it. Such a load-dependent adhesion-failure mechanism has been proposed theoretically^70,71^, providing a plausible route by which elevated contractility destabilizes basal attachment and promotes cell detachment and extrusion. Depending on matrix properties, increased basal contractility and adhesion may either promote cell–ECM detachment when adhesions fail under excessive load, or favor basal invasion when the ECM is sufficiently deformable and remodelable, as in collagen gels^11^. Interestingly, basally-oriented extrusions were also recently observed in 3D organoids of hypercontractile mouse tumor cells as well as MCF10A, MDCK cell cysts and in 2D intestinal organoids^11,46,72^, emphasizing the physio-pathological implication of the mechanism we describe here.

Strikingly, RhoA-induced basal extrusions were largely caspase-negative. While apical extrusion is typically associated with apoptotic elimination, basal extrusion may represent a survival route for hypercontractile cells^14,73,74^. Furthermore, RhoA-activated cells exhibited reduced E-cadherin expression and cell-cell adhesion. This phenotype may facilitate basal extrusion and potential ECM invasion, as shown in a previous study^11^. We also observed that basally-oriented cell extrusions are independent of both caspase activation and the well-studied stretch-activated ion channel Piezo-1. These observations might appear to conflict with studies that have shown Piezo-1 requirement for live cell extrusion in cultured epithelial monolayers^2,55,75,76^, zebrafish and mouse^8^. One explanation is that the Piezo-1 activation and downstream signaling pathways are required for cell extrusion under compressive stress, as opposed to the tensile stress observed in this study. This suggests a non-canonical pathway for cells undergoing extreme values of tension, as has been proposed in previous works^7,28,77^. Altogether, our work supports a model in which hypercontractile cells are eliminated through a RhoA-dependent, tension-sensitive pathway that bypasses canonical apoptotic signaling. Crucially, extrusion orientation dictates fate when cells are grown on a soft and permissive ECM: while apical extrusions are mostly apoptotic, basal extrusions are initially caspase-independent and may facilitate survival and invasion. This coupling between mechanics, extrusion route, and fate highlights a previously unrecognized mechanism of epithelial regulation with broad implications for development, tissue homeostasis, and cancer progression.

## Resource Availability

### Lead contact

Further information and requests for resources and reagents should be directed to and will be fulfilled by the Lead Contact, Simon de Beco.

### Materials availability

The cell lines used in this paper are available from the Lead Contact upon request.

### Data and code availability

The raw imaging datasets and codes for mechanical stress analysis have not been deposited in a public repository, but are available from the Lead Contact upon request.

## Acknowledgements

We thank the members of the ‘Cell Adhesion and Mechanics’ team (Institut Jacques Monod) as well as Drs Nicolas Minc and Romain Levayer for helpful discussions. We acknowledge the ImagoSeine core facility of the Institut Jacques Monod, member of France-BioImaging (ANR-10-INBS-0004) and IBiSA, with support of Labex ‘Who Am I’, Inserm Plan Cancer, Region Ile-de-France and Fondation Bettencourt-Schueller. We thank Xavier Trepat for sharing the MDCK Opto-RhoA cell line. This work was supported by the European Research Council (grant no. Adv-101019835 “DeadorAlive” to B.L.), Labex ‘Who Am I’ (ANR-11-LABX-0071 to F.W., B.L., R.-M.M. and S.d.B.), the Alexander von Humboldt Foundation (Alexander von Humboldt Professorship to B.L.), the Ligue Contre le Cancer (Equipe labellisée 2019 to R.-M.M. and B.L.), the CNRS through 80|Prime program (to B.L.), Institut National du Cancer INCa (‘Invadocad’, PLBIO18-236) and the Agence Nationale de la Recherche (“STRATEPI” DFG-ANR-22-CE92-0048 and “VISCOMAG2” ANR-24-CE42-6142 to RM.M. and B.L.). F.W. received funding from the European Research Council and Labex ‘Who Am I’. L.A. received funding from the Ligue contre le Cancer. Mallica Pandya was supported by a studentship from the CDT in Biodesign Engineering funded by the Engineering and Physical Sciences Research Council (EPSRC). GC was supported by grant BB/W011123/1 from the Biotechnology and Biological Sciences Research Council. A.D. acknowledges funding from the Novo Nordisk Foundation (grant no. NNF18SA0035142 and NERD grant no. NNF21OC0068687), Villum Fonden (grant no. 29476) and the European Union (ERC, PhysCoMeT, 101041418). Views and opinions expressed are however those of the authors only and do not necessarily reflect those of the European Union or the European Research Council. Neither the European Union nor the granting authority can be held responsible for them.

## Author Contributions

A.D., B.L. and S.D.B conceived the study. F.W., P.G., R.D., L.A., A.S., M.A., T.D. performed experiments and subsequent analyses. T.M., S.M. performed numerical simulations and subsequent analyses. M.P. and G.C provided resources. F.W., T.M., R.M.M., A.D., B.L. and S.D.B wrote the manuscript.

## STAR Methods

KEY RESOURCES TABLE.

| REAGENT or RESOURCE | SOURCE | IDENTIFIER |
|---|---|---|
| Antibodies |  |  |
| Anti-cleaved caspase-3 (1:100) | Sigma Aldrich | Cat# C8487 |
| Anti-phospho-myosin light chain 2 (Ser19) (1:100) | Cell Signaling Technology | Cat# 3671 |
| Anti-E-cadherin, rabbit mAb, clone 24E10 (1:100) | Cell Signaling Technology | Cat# 3195 |
| Anti-paxillin, rabbit mAb, clone Y113 (1:100) | Abcam | Cat# ab32084 |
| Anti-β-catenin (1:200) | Sigma Aldrich | Cat# C2206 |
| Anti-α-catenin (1:200) | Sigma Aldrich | Cat# C2081 |
| Bacterial and virus strains |  |  |
| Biological samples |  |  |
| Chemicals, peptides, and recombinant proteins |  |  |
| DMEM, high glucose, GlutaMAX, pyruvate | Life Technologies | Cat# 10569010 |
| Fetal bovine serum (FBS) | Invitrogen | Cat# A5256701 |
| Penicillin-Streptomycin | Invitrogen | Cat# 15140122 |
| Trypsin-EDTA 0.05% | Merck | Cat# 25200056 |
| ML-7 | Cellchem | CAS 109376-83-2 |
| Mitomycin C | Sigma Aldrich | Cat# M4287 |
| Rho Activator II (CN03) | Cytoskeleton Inc. | Cat# CN03 |
| SPY-DNA 650nm | Spirochrome | Cat # CY-SC501 |
| SPY-FastActin 650nm | Spirochrome | Cat # CY-SC505 |
| Annexin V, Alexa Fluor 647 conjugate | Invitrogen | Cat# A23204 |
| Z-VAD-FMK (caspase inhibitor) | Promega | Cat# G7231 |
| Fibronectin, bovine | Sigma Aldrich | Cat# F8630-16 |
| Rat tail collagen I | Corning | Cat# CLS354236 |
| HEPES 1 M | Invitrogen | Cat# 15630080 |
| Opti-MEM I Reduced Serum Medium, 100 mL | Invitrogen | Cat# 31985062 |
| PDMS CY52-276A / CY52-276B | Dow Corning Toray | CY52-276A; CY52-276B |
| (3-Aminopropyl)trimethoxysilane | Sigma Aldrich | Cat# 281778 |
| 3-(Trimethoxysilyl)propyl methacrylate | Sigma Aldrich | Cat# 440159 |
| Acrylamide | BIORAD | Cat# 1610140 |
| Bis-acrylamide | BIORAD | Cat# 1610142 |
| Ammonium persulfate | Sigma Aldrich | Cat# 1610700 |
| TEMED | Thermo Fischer Scientific | Cat# 17919 |
| Sulfo-SANPAH | Sigma Aldrich | Cat# 803339 |
| Sodium dodecyl sulfate (SDS) | EuroMedex | Cat# EU0660 |
| Paraformaldehyde | Thermo Fisher Scientific | Cat# 441244 |
| Triton X-100 | Sigma Aldrich | CAS:9002-93-1 |
| Bovine serum albumin (BSA) | Sigma Aldrich | Cat# A8022 |
| Hoechst | Thermo Fisher Scientific | Cat# H3570 |
| Alexa Fluor 647 Phalloidin (1:250) | Life Technologies | Cat# A22287 |
| ProLong Gold Antifade Mountant | Thermo Fisher Scientific | Cat# P36961 |
| Critical commercial assays |  |  |
| Deposited data |  |  |
| Experimental models: Cell lines |  |  |
| MDCK WT Type II | ATCC | ECACC 00062107 |
| MDCK Cry2mCherry;CAAX-CIBN_GFP (Opto-Ctrl) | Laboratory of Benoit Ladoux | N/A |
| MDCK Cry2mCherry-ARHGEF11;CAAX-CIBN-GFP (Opto-RhoA) | Laboratory of Xavier Trepat | N/A |
| MDCK Cry2mCherry-Tiam1; CAAX-CIBN-GFP (Opto-Rac1) | Laboratory of Benoit Ladoux | N/A |
| MDCK Cry2mCherry-ARHGEF11;N-Shroom-CIBN-GFP (Shroom-RhoA) | Laboratory of Guillaume Charras | N/A |
| MDCK Cry22mCherry-ARHGEF11;ITB1-mVenus-CIBN (ITB1-RhoA) | Laboratory of Guillaume Charras | N/A |
| Experimental models: Organisms/strains |  |  |
| Oligonucleotides |  |  |
| Recombinant DNA |  |  |
| Software and algorithms |  |  |
| MATLAB | MathWorks | 2022b |
| Fiji | https://imagej.net | v1.54q |
| Image Stabilizer plugin (ImageJ) | https://imagej.net/plugins/image-stabilizer | N/A |
| OrientationJ plugin (Fiji) | Biomedical Imaging Group, EPFL | N/A |
| PIVlab v3.08 | Pivlab.fr | v3.08 |
| BISM (Bayesian inversion stress microscopy) | Nier et al.<sup>26</sup> | N/A |
| FSegment | Rogge et al.<sup>80</sup> | N/A |
| Python | Python Software Foundation | N/A |
| Custom Python code for 3D PIV and 3D FTTC (2.5D TFM) | This paper; adapted from Böhringer et al.<sup>78</sup> and Blumberg & Schwarz<sup>79</sup> | N/A |
| DataViewer V2 | Optics11 Life | N/A |
| Other |  |  |
| Glass-bottom dishes, 35 mm (FluoroDish) | World Precision Instruments (WPI) | Cat# FD35-100 |
| FluoSpheres fluorescent beads, 647 nm (2D/2.5D TFM) | Invitrogen | Cat# F8807 |
| FRAP module iLas2 | Gataca Systems | N/A |
| Chiaro nanoindenter | Optics11 Life | N/A |

## EXPERIMENTAL MODEL AND STUDY PARTICIPANT DETAILS

### Cell culture and cell lines

MDCK WT (MDCK WT), MDCK Cry2-mCherry;CAAX-CIBN_GFP (Opto-Ctrl), MDCK Cry2-mCherry-ARHGEF11;CAAX-CIBN-GFP (Opto-RhoA), MDCK Cry2-mCherry-Tiam1;CAAX-CIBN-GFP (Opto-Rac1), Cry2-mCherry-ARHGEF11;ITB1-CIBN-GFP (ITB1-RhoA) and Cry2-mCherry-ARHGEF11;N-Shroom-CIBN-GFP (Shroom-RhoA) are MDCK strain II stable cell lines that were maintained in Dulbecco’s modified Eagle’s medium composed of high glucose and pyruvate (DMEM GlutaMAX, Life Technologies), supplemented with 10 % Fetal bovine serum (FBS, Invitrogen), 100µg/ml Penicillin, 100µg/ml Streptomycin (Invitrogen). Cells were passaged every 3 days using 0.05% Trypsin-EDTA (Merck) after being washed in 1X PBS. Cells were incubated at 37°C in 5% CO2 unless otherwise described. Cells were seeded 24 hours before the experiments to reach confluency the day of the experiment.

MDCK Opto-RhoA were a kind gift by Xavier Trepat (IBEC, Barcelona). Plasmids encoding Cry2-mCherry-Tiam1 and Cry2-mCherry were provided by Mathieu Coppey (Institut Curie, Paris). Plasmids encoding N-Shroom-CIBN-GFP and CIBN-GFP were obtained from Addgene (plasmids #170976, #26867). Integrin Beta1-mVenus (ITB1-mVenus) was a kind gift from Dr. Joan Brugge (Harvard Medical School, Boston, MA, USA). Lentiviral backbones carrying an IRES-Puro cassette were designed and generously provided by Dr. Stephen Terry (UCL). ITB1-mVenus was fused to CIBN using standard cloning approaches. N-Shroom-CIBN-GFP and ITB1-mVenus-CIBN were inserted into the lentiviral backbones using standard approaches.

Lentiviruses carrying Cry2-mCherry-ARHGEF11, CIBN-GFP-CAAX, N-Shroom-CIBN-GFP, ITB1-mVenus-CIBN, Cry2-mCherry-Tiam1 and Cry2-mCherry were generated by transfection into HEK293T cells using Lipofectamine 3000 (Thermo Scientific #L3000008). Wild-type MDCK II cells were transduced with lentiviruses and selected with puromycin (1 µg/mL; InvivoGen). After selection, cells were sorted by flow cytometry, retaining the brightest 1% of m-Cherry and GFP-positive cells.

## METHOD DETAILS

### Reagents and treatments

To reduce actomyosin contractility, cells were treated with 30 µM ML-7 (Cellchem), added to the culture medium 1 hour prior to imaging. To inhibit cell proliferation, cells were incubated with 10 µg/ml Mitomycin-C (Sigma-Aldrich) for 1 hour; the medium was then replaced with fresh, warm medium, and cells were imaged 8 hours after the Mitomycin-C treatment. To activate RhoA in MDCK WT cells, CN03 (Cytoskeleton Inc.) was added at 1.5 µg/ml for 8 hours. For live imaging of fluorescently labeled cell nuclei and actin, confluent MDCK monolayers were incubated with either SpyDNA or SpyActin FAST (Spirochrome) at a 1:1000 dilution, 3 hours before the start of the experiment, and maintained in the medium throughout. To monitor apoptosis, Annexin V (Abcam) was added at a 1:1000 dilution 1 hour prior to imaging and maintained for the duration of the experiment. To inhibit caspase activation, 50 µM Z-VAD-FMK (Promega) was added to 2 ml of medium 1 hour before the experiment and maintained throughout the imaging period.

### Co-culture experiments

MDCK WT and MDCK Opto-RhoA cells were resuspended separately in 3 mL of pre-warmed culture medium. Cells were then mixed at the desired ratio (3% or 50% of Opto-RhoA) and seeded onto glass-bottom dishes coated with 50 µg/mL fibronectin, with a final seeding density of 1.2 × 10^6 cells in total. After 24 h of culture, cells were rinsed once with 1× PBS and fresh pre-warmed medium was added prior to imaging.

### Collagen Matrix Experiments

We mixed Rat tail collagen 1 (Corning) with H2O, Hepes 1M, NaOH 0.34M and MEM 1X to reach a final concentration of 2.5 mg/ml of collagen. We deposited 150µl of the mix in glass bottom dishes (Fluorodish, WPI), and incubated for polymerization at 37°C with 5% CO2 for 1 hour. 2 ml of warm media were added to the polymerized collagen matrix. 8.10^5 cells were mixed in 2ml of media then seeded in the dish with the matrix 24 hours before the start of the experiment.

### Traction Force Microscopy

All PDMS substrates were made in 35 mm diameter glass-bottom dish (Fluorodish). CY52-276A and CY52-276B polydimethylsiloxane (PDMS; Dow Corning Toray) were mixed in a weight ratio of 1:1 (15 kPa) or 6:5 (2kPa). 0.1 g PDMS drops were deposited in the center 35 mm diameter glass-bottom dish (Fluorodish, WPI), spin-coated (30s at 500 rpm, acceleration 100rpm/sec), and cured for 2h at 80°C. The substrates were then functionalized with human fibronectin (50 μg/ml) incubated on the substrate for 1 hr prior to cell seeding.

For 2D TFM, CY52-276A and CY52-276B polydimethylsiloxane (PDMS; Dow Corning Toray) were mixed in a weight ratio of 1:1 (15 kPa). 0.1 g PDMS drops were deposited in the center 35 mm diameter glass-bottom dish (Fluorodish, WPI), spin-coated (30s at 500 rpm, acceleration 100rpm/sec), and cured for 2h at 80°C. The substrates were then functionalized with a 10% (3-Aminopropyl) trimethoxysilane (Sigma-Aldrich) in ethanol for 15 minutes, rinsed twice with absolute ethanol and dried for 1 minute at 80°C. 0.25 µm 647nm carboxylate fluorescent beads (Invitrogen) were filtered with a 0.45µm filter and functionalized on the substrate at 4:1000 dilution in H2O. Pure human fibronectin (50 μg/ml) was incubated on the substrate for 1 hr prior to cell seeding. Between each step, the samples were rinsed 3 times with PBS.

For 2.5D TFM, 30kPa soft polyacrylamide (PAA) gels were used. Briefly, Fluorodishes (Fluorodish, WPI) were incubated for 30 min in a silane solution containing 0.07% (trimethoxysilyl)propylmethacrylate (Sigma-Aldrich, no. 440159) and 0.07% acetic acid in ethanol. The silanized fluorodishes were rinsed with ethanol, dried by evaporation, and stored at room temperature. A freshly prepared polyacrylamide gel mixture (acrylamide, bis-acrylamide, ammonium persulphate, and TEMED in PBS, containing 1% 647 nm 0.22 nm fluorescent beads (FluoSpheres, Invitrogen)) was cast onto the coverslips and allowed to polymerize at room temperature for 1h to form a ∼ 100-µm-thick gel. The resulting substrates were stored in PBS at 4 °C. Before use, PAA gels were functionalized with Sulpho-SANPAH (Cultek), activated under 365-nm UV light for 7min30, and coated with a solution of bovine fibronectin (50 µg/ml, Sigma-Aldrich) in PBS, followed by overnight incubation at 4 °C. Prior to cell seeding, gels were rinsed three times with PBS. Twenty-four hours before imaging, 0.6 × 10⁵ cells were resuspended in warm culture medium and seeded onto the PAA gels.

### Immunostaining and antibodies

Samples were fixed with 4% paraformaldehyde (Thermo Fisher Scientific, Waltham, MA, USA) for 10 mins and permeabilized with 0.5% Triton-X for 5 mins. Cells were then blocked with 1% BSA/PBS and incubated with primary antibody overnight with agitation at 4°C. Cells were incubated in secondary antibody for 3 hours, washed and incubated with Hoechst (Thermofisher scientific) 1:10000 for 10 mins and then mounted on another glass coverslip using ProLong Gold Antifade Mountant (ThermoFisher Scientific). For immunostaining, the following primary antibodies were used: Caspase-3 cleaved (C8487 Sigma-Aldrich (St Louis, MO)) 1:100, Phospho-myosin II P-Ser19 (3671S Cell Signaling Technology (Danvers, MA)) 1:100, E-cadherin (24E10-Cell Signaling Technology) 1:100, Paxillin (Abcam, Ab32084) 1:100, Beta-catenin (C2206 Sigma-Aldrich (St Louis, MO)) 1:200, alpha-catenin (C-2081 Sigma-Aldrich (St Louis, MO)) 1:200. The following secondary antibodies were used, Anti-rabbit (A31573 Life Technologies Ltd. (Paisley, UK)) 1:250 and anti-mouse (A31571 Life Technologies Ltd. (Paisley, UK)) 1:250. Phalloidin (A22287 Life Technologies Ltd. (Paisley, UK)), 1:250.

#### Microscopic imaging of live and fixed samples

Live imaging for 2D TFM experiments was performed with a Zeiss spinning disk confocal (CSU-X1) using a 25x oil objective. Local illumination for restricted optogenetic activation was achieved with a 488 nm laser controlled by a FRAP module (ILaS2, Gataca systems). Cells and fluorescent beads were imaged with a frame rate of 0.5 frame/min for 10 hours with a Z range of 6µm, interval 3µm. Live imaging for 2.5D TFM experiments was performed using a Nikon spinning-disk confocal microscope (CSU-W1) equipped with a 25× air objective. Cells were photostimulated at a single focal plane using a 488 nm laser every 7 min. Fluorescent bead images were acquired at the same temporal resolution (1frame/7min) with Z-stacks spanning 41 µm and acquired with a 1 µm step size, for a total duration of 12 h. Live imaging of confluent monolayers on collagen-1 matrices was performed with a Nikon spinning disk confocal microscope (CSU W1) using a 25X air objective, with a frame rate of 0.5 frame/min for 12 hours with a Z-range of 20 µm, interval between 0.5 and 1 µm. For high resolution images, a Confocal LSM 980 Airyscan (Zeiss) was used with 63X and 100X oil immersion objectives.

### Laser ablation

After optogenetic activation for 10 minutes, laser ablations were done by illuminating a rectangle of 150 µm x 45 µm with an ultra-violet 355nm laser at high power using a FRAP module (ILaS2, Gataca systems) on a Zeiss spinning disk confocal (CSU X1). Cells were then imaged using a 568nm laser (mCherry channel) every 5 sec for 5 mins. The recoil velocity was measured by manually detecting cell edges over time and plotting the distance from their initial position.

### Cell stiffness measurement

We used an indenter (Chiaro nanoindenter, Optics11 Life) with a soft probe (k = 0.015 N m–1; tip radius, 3 μm) to measure cell stiffness. MDCK Opto-RhoA cells were grown until confluency on fibronectin-coated glass substrates. Measurements were performed on monolayers after 10 minutes of continuous optogenetic stimulation at 488nm, or without any stimulation. Each measurement was done with an indentation depth of 1.5µm during 4 seconds. To determine the elastic modulus, the loading curve was analyzed using the built-in software (DataViewer V2, Optics11 Life). The analysis is based on the Hertz model (Hertzian contact), which assumes a linear elastic response of the sample. The single-fit method was used with a maximum load (Pmax) of 90% and a Poisson’s ratio of 0.5.

## QUANTIFICATION AND STATISTICAL ANALYSIS

### 2D TFM analysis

The bead images obtained during TFM experiments were merged with the corresponding reference bead images taken after sodium dodecyl sulfate (SDS) treatment. The resulting stack of images was preprocessed using the Image Stabilizer plug-in in ImageJ and the illumination was corrected to remove background noise. Displacement field of beads was obtained using PIVlab (v. 3.08), a particle image velocimetry toolbox developed in MATLAB, with an interrogation window of 32 × 32 pixels and an overlap of 50%. Bead displacements were then correlated to a traction force field using Fourier transform traction cytometry (FTTC), a known theoretical substrate stiffness and a regularization parameter of 9 × 10 . From the traction force field, we were able to infer the stress tensor everywhere in the tissue using BISM with a regularization parameter of Λ = 10^−6^. Isotropic stress was calculated as half the trace of the stress tensor. To generate the heat map of isotropic stress and traction force magnitude, smoothing was applied through linear interpolation.

### 2.5D#TFM analysis

Raw 3D bead volumes were sampled to isotropic voxels (0.32 µm) via tricubic interpolation. Three-dimensional substrate displacements were measured by 3D PIV between each time-frame bead volume and the reference stack using a custom python implementation in part using logic and code from Böhringer et al^78^.PIV interrogation windows of 6×6×6 µm were used with 67% overlap. Spurious vectors were removed and replaced by nearest-neighbor interpolation. A drift correction was applied to the displacement field. Substrate tractions were recovered from the displacement field (v_x_(x, y), v_y_(x, y), v_z_(x, y)) at a top layer of the substrate by FTTC extended to three components (f_x_, f_y_, f_z_). A finite-thickness Boussinesq Green’s function accounting for the rigid glass substrate at depth h =200 µm was used^45^. The forward problem relates tractions to displacements via this Green’s tensor in Fourier space; the inverse problem was solved with Tikhonov regularization and the optimal regularization parameter was determined per image by generalized cross-validation using logic and code from Blumberg & Schwarz^79^. Positive f_z_ indicates outward displacement (toward the cells); negative fz indicates compression into the substrate.

#### Extrusion rate quantification

Extrusions were tracked manually and the extrusion rate was measured by dividing the number of cell extrusion events by the area of the stimulated or non-stimulated area, normalized by the total number of extrusions in the field of view. To assess the spatial distribution of cell extrusion events, we transformed the imaging field to a distance map and matched each extrusion’s coordinates to its corresponding zone. We then quantified the number of extrusion events per zone and normalized this count by the total number of extrusions. To correct for differences in zone sizes, we calculated the proportion of the field area occupied by each zone (in pixels) and adjusted the raw extrusion frequencies accordingly. The final extrusion probability per zone was obtained by normalizing the size-adjusted frequencies across all zones.

#### Stress Fibers segmentation

Stress fibers were quantified and segmented using the Fsegment application^80^ on Matlab. Images of actin stress fibers were preprocessed in the application with a global threshold of 0.1% of maximum intensity, and a top-hat transformation with a size of element of 3 pixels. Only thin fibers with a size of 1-to-5 pixel width and 3 pixel length minimum were detected with a step for the algorithm of 5 pixels. The binary masks corresponding to the stress fibers for each frame of time were saved and used afterward. Stress fibers orientation was measured using the OrientationJ plugin on Fiji software, with a local window of 4 pixels with Hessian method and HSV colormap display. The average orientation was obtained by extracting orientation values in degrees from the orientation matrix with a binary mask of the cell. The mean order parameter of the corresponding orientation was computed from the extracted values of orientation.

### Statistical analysis

All statistical tests have been run using the Matlab program.

All the statistical details of experiments can be found in the figure legends.

In all descriptions, N represents the numners of independent experiments, n represents the number of events (extrusions, cells…).

In all plots, the meaning of center and dispersion have been precised in figure legends.

## Supplemental videos

**Video S1: Optogenetic recruitment of CRY2-mCherry and ArhGEF11-CRY2-mCherry to the plasma membrane after light stimulation. Related to Figure 1B**

Time lapse movies of the mCherry signal of MDCK Opto-Ctrl **(left)** and MDCK Opto-RhoA cells **(right)** optogenetically stimulated within a region of 150 x 150µm (white square). Scale bar: 50µm.

**Video S2: Examples of cell extrusion. Related to Figure 2A**

Time lapse movies showing cell extrusions (white arrowheads) of MDCK Opto-RhoA **(left)** and MDCK Opto-Ctrl **(right)** optogenetically stimulated within a region of 150×150µm (white square). Scale bar: 50µm.

**Video S3: Optogenetic recruitment of ArhGEF11-CRY2-mCherry to the plasma membrane after light stimulation in regions of various sizes and shapes. Related to Figure 4A**

Time lapse movies of the mCherry signal for the Opto-RhoA cell line during stimulation at 488nm in regions of varying sizes: 44 x 44 µm **(left)**, 50 x 50 x 50 µm **(middle),** 260 x 260µm **(right).** Scale bar: 50µm.

**Video S4: Alignment of actin stress fibers prior to cell extrusion. Related to Figure 5F**

Time lapse movies showing actin dynamics (stained with FAST-Act, Spirochrome) in MDCK Opto-RhoA and Opto-Control cells cultivated on glass. Extruding cells are shown with an asterisk. Actin is shown at the apical junction **(left)** and basal plane **(right)**. Movies start 5 hours prior to extrusion onset. Magnification 63X. Frame rate: 0.5/min. Scale bar: 10µm.

### Video S5: Delayed appearance of Annexin V signal after extrusion onset. Related to Figure 6E

Time lapse movies of CIBN-CAAX-GFP (membranes) and Annexin V signals showing apical (top) and basal (bottom) extrusions of blue-light-illuminated MDCK Opto-RhoA cells on Collagen 1 matrices. **Left:** xy view**, right:** ZY view. Time t = 0 min corresponds to the onset of extrusion. Annexin V signal detection is represented with a white arrowhead. Scale bar: 10µm.

