## Supplemental Information for "Mechanical Tension Actively Triggers RhoA-Mediated Cell Extrusion"

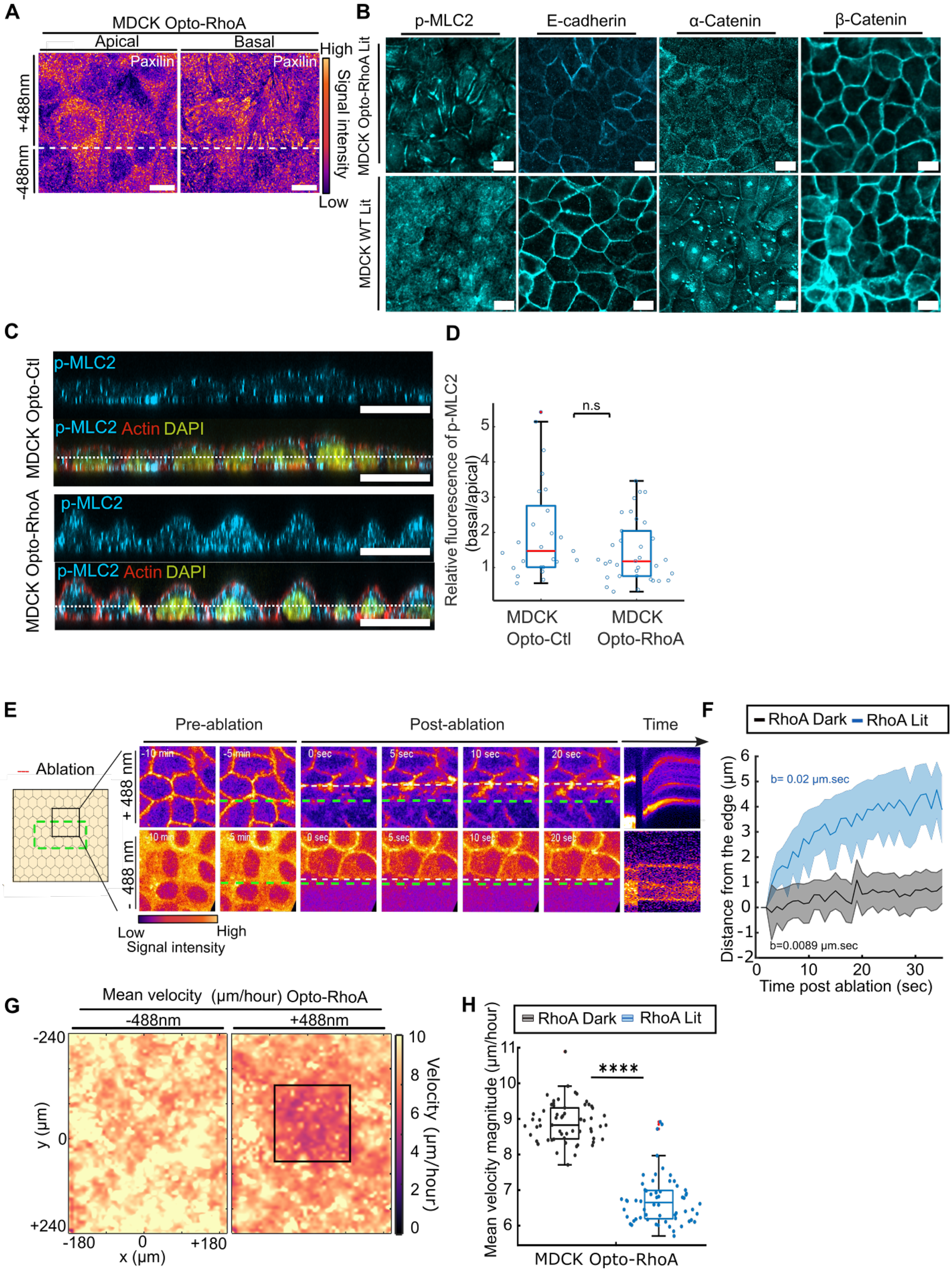

A
MDCK Opto-RhoA
Apical
Basal
Paxilin
Paxilin
+488nm
-488nm
High
Signal intensity
Low
B
p-MLC2
E-cadherin
α-Catenin
β-Catenin
MDCK Opto-RhoA Lit
MDCK WT Lit
C
MDCK Opto-Ctl
MDCK Opto-RhoA
p-MLC2
p-MLC2 Actin DAPI
D
Relative fluorescence of p-MLC2 (basal/apical)
n.s
MDCK Opto-Ctl
MDCK Opto-RhoA
E
Ablation
Pre-ablation
Post-ablation
Time
+ 488 nm
- 488 nm
-10 min
-5 min
0 sec
5 sec
10 sec
20 sec
Low
High
Signal intensity
F
RhoA Dark
RhoA Lit
b= 0.02 µm.sec
b=0.0089 µm.sec
Distance from the edge (µm)
Time post ablation (sec)
G
Mean velocity (µm/hour) Opto-RhoA
-488nm
+488nm
y (µm)
x (µm)
Velocity (µm/hour)
H
RhoA Dark
RhoA Lit
****
Mean velocity magnitude (µm/hour)
MDCK Opto-RhoA

**Figure S1: RhoA-induced cell contractility affects cell junctions and intercellular stress. Related to Figure 1.**

**A** MDCK Opto-RhoA cells cultured on glass coverslips until confluency and stimulated at 488 nm within a restricted square area, then fixed and stained for paxillin. Apical views (left) and basal views (right) are shown. The white dashed line indicates the boundary between the stimulated (+488 nm) and unstimulated (−488 nm) regions. Scale bar = 10µm. **B** MDCK Opto-RhoA and MDCK WT cells cultured until confluency on glass substrate then globally stimulated at 488 nm (Lit), fixed and stained for p-MLC2, E-cadherin, alpha-catenin, beta-catenin. Scale bar=10µm. **C** Representative orthogonal confocal sections of MDCK Opto-Ctl and MDCK Opto-RhoA monolayers, globally stimulated and immunostained for phosphorylated myosin light chain 2 (p-MLC2, cyan), F-actin (red) and nuclei (DAPI, yellow). The dashed line indicates the boundary between the apical and basal side of the epithelial monolayer and was used as a reference for p-MLC2 fluorescence intensity measurements. Scale bars, 20 µm. **D** Relative p-MLC2 fluorescence intensity ratio between the basal and apical sides in stimulated MDCK Opto-Ctl (N=1 n=24 cells) and MDCK Opto-RhoA cells (N=1 n=33 cells). The p-MLC2 intensity ratio was calculated by dividing the mean fluorescence intensity measured at the basal side of each cell by the corresponding apical intensity. Wilcoxon rank-sum test: n.s., $p = 0.1056$. **E** Time-lapse images following laser ablation. The green dashed lines indicate the ablated region, while the white dashed lines highlight cell displacement after the cut for the MDCK Opto-RhoA that were globally stimulated for 20 minutes prior to ablation (+488nm) or not stimulated (-488nm). **F** Quantification of the recoil velocity following laser ablation in MDCK Opto-RhoA monolayers stimulated at 488 nm (RhoA Lit) (N = 2 n = 11) or maintained under unstimulated conditions (RhoA Dark) (N = 2 n = 10). Cell displacement was measured by tracking membrane movement after ablation. **G** Mean velocity magnitude maps of MDCK Opto-RhoA monolayers after 1 hour of restricted stimulation within a 150 × 150 µm square area, compared with the unstimulated condition before activation (left). Cell velocities were quantified using particle image velocimetry (PIV) analysis and analyzed in Matlab. **H** Distribution of mean velocity values shown in **(G).** Median values of cell velocity in the stimulated (RhoA Lit) and unstimulated cells (RhoA Dark). (N=3 n= 40). Wilcoxon rank-sum test: **** $p = 2.51 \times 10^{-19}$. Curves represent the mean +/- s.e.m. Box plots represent the median, interquartile (box), and 1.5 IQR (whiskers).

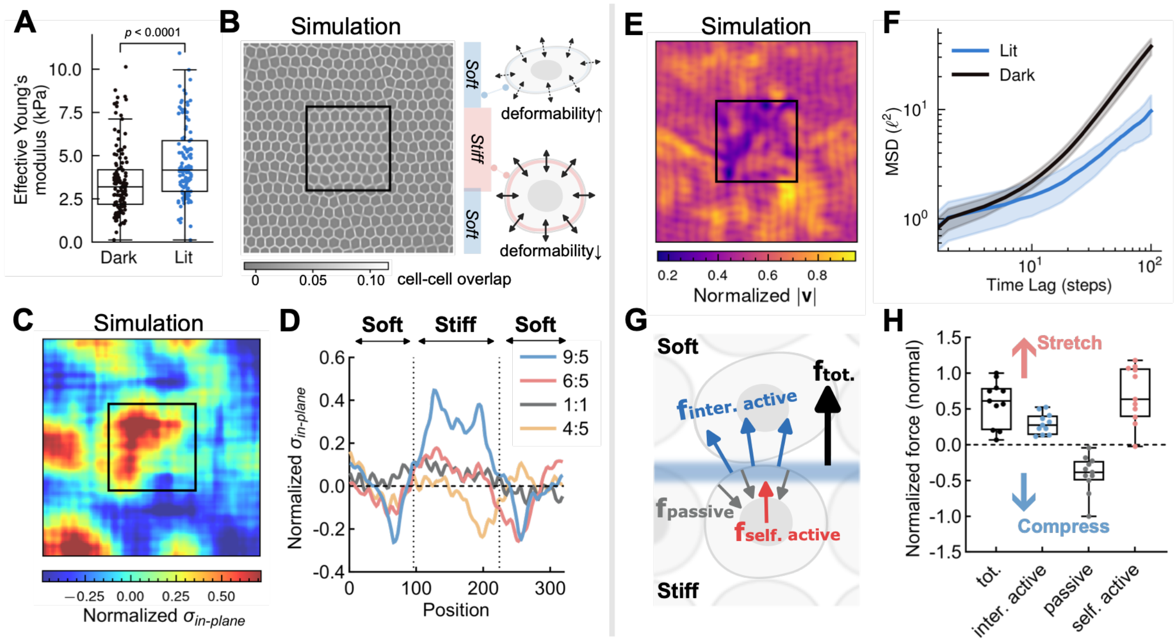


**Figure S2: RhoA-induced contractility elevates cellular stiffness and drives in-plane tension buildup via mechanical asymmetry. Related to Figure 1.**

**A** Atomic force microscopy (AFM) measurements reveal a significant increase in effective Young's modulus in RhoA-activated (Lit) cells, confirming contractility-induced stiffening (****$P < 0.0001$, Mann–Whitney test). **B** Schematic of the simulated monolayer with a central region of increased stiffness, mimicking experimental RhoA activation. Colormap shows cell-cell phase overlap, calculated by $\Sigma_{i<j}\Phi_i\Phi_j$. **C** Time-averaged map of normalized in-plane isotropic stress ($\sigma_{in\text{-}plane}$) shows localized tension buildup in the activated region. **D** Line profiles of $\sigma_{in\text{-}plane}$ under varying stiffness ratios (stiff:soft) demonstrate that tension amplitude increases with stiffness mismatch. Each curve represents the mean value across 3 independent simulations. **E** Simulated time-averaged map of normalized velocity magnitude shows locally suppressed motility within the activated region. **F** Simulated mean-square displacements comparing cells inside the activated region (Lit) and outside (Dark), demonstrating reduced cell motion in the Lit region. $\ell$ denotes the simulation spatial unit (3 independent simulations; mean ± SD). **G** Schematic illustrating the decomposition of total forces acting at the interface between activated and non-activated regions into passive (gray), intercellular active (blue), and self-propulsion (red) components. **H** Quantification of force density components normal to the interface reveals that while passive forces are inward-directed (compressive), both active intercellular and self-propulsion forces point outward, together driving the buildup of in-plane tension in the activated region (3 simulations, each point represents the result for one square edge in each simulation). Box plots represent the median (center line), interquartile range (box), and minimum to maximum values (whiskers).

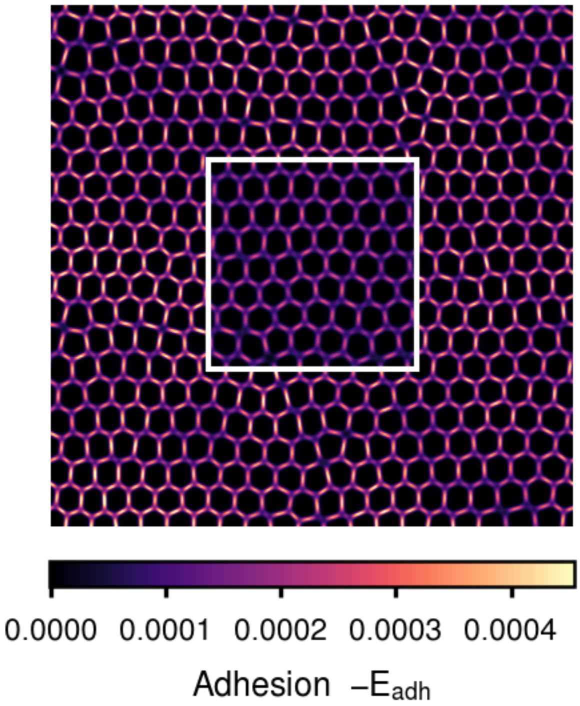


**Figure S3: Cell-cell adhesion in the simulated monolayer. Related to Figure 2.**
$-E_{adh}$ represents the local adhesion strength[S1], where larger positive values indicate stronger adhesive interactions. The white square marks the activated region.

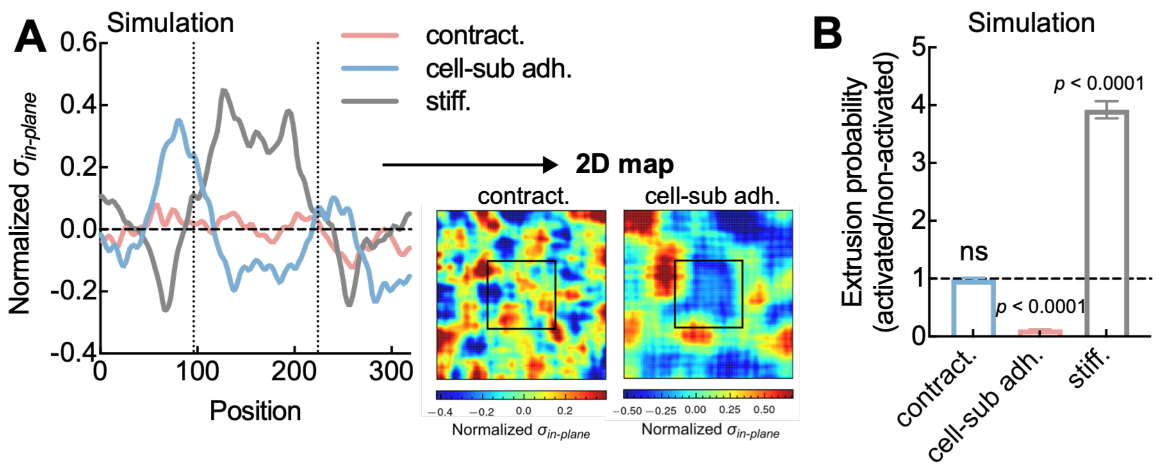


**Figure S4: Neither increased contractility nor increased cell–substrate adhesion alone reproduces the experimentally observed tension buildup and elevated extrusion. Related to Figure 2.**

**A** Line profiles of normalized in-plane isotropic stress ($\sigma_{\text{in-plane}}$) across the monolayer for three conditions: increased basal contractility in the activated region ($\zeta_Q = -2\zeta_S$; contract., red), increased cell–substrate adhesion in the activated region (2× the non-activated value; cell-sub adh., blue), and increased stiffness in the activated region (9:5 ratio, corresponding to the data in Fig. S2D; stiff., gray). Dashed vertical lines mark the boundaries of the activated region. **Insets:** time-averaged 2D maps of $\sigma_{\text{in-plane}}$ for the contract. and cell-sub adh. conditions; the stiff. map is shown in Fig. S2C. Enhanced contractility alone produces no appreciable change in $\boldsymbol{\sigma_{\text{in-plane}}}$ within the activated region, whereas increased cell–substrate adhesion reduces $\boldsymbol{\sigma_{\text{in-plane}}}$ there — neither recapitulates the tension buildup observed experimentally and reproduced by stiffness modulation. **B** Relative extrusion probability (activated/non-activated) for the same three conditions. Increased contractility alone does not significantly elevate extrusion in the activated region, and increased cell–substrate adhesion suppresses it. Data are mean ± SEM from 3 independent simulations; one-sample t-test against the null value of 1 (ns, not significant; ****P < 0.0001).

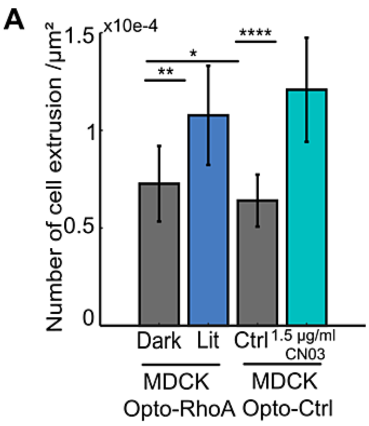


**Figure S5: RhoA-induced tension is sufficient to induce cell extrusion. Related to Figure 2.**
**A** Quantification of cell extrusion events in MDCK Opto-RhoA and MDCK Opto-Ctrl monolayers grown on collagen I matrices. MDCK Opto-RhoA cells were globally stimulated for 8 hours (Lit) (N=2, n=304 cells) or maintained under unstimulated conditions (Dark) (N=2, n=125 cells). MDCK Opto-Ctrl cells were treated with 1.5 µg/mL CN03 (N=2, n=347 cells) or left untreated (Ctrl) (N = 2, n =184 cells). Cells were fixed and stained with DAPI. Bar graphs represent mean ± s.e.m. Wilcoxon rank-sum test: ;** p = 0.01744; * p = 0.03402; **** p = $5.11 \times 10^{-8}$, from left to right.

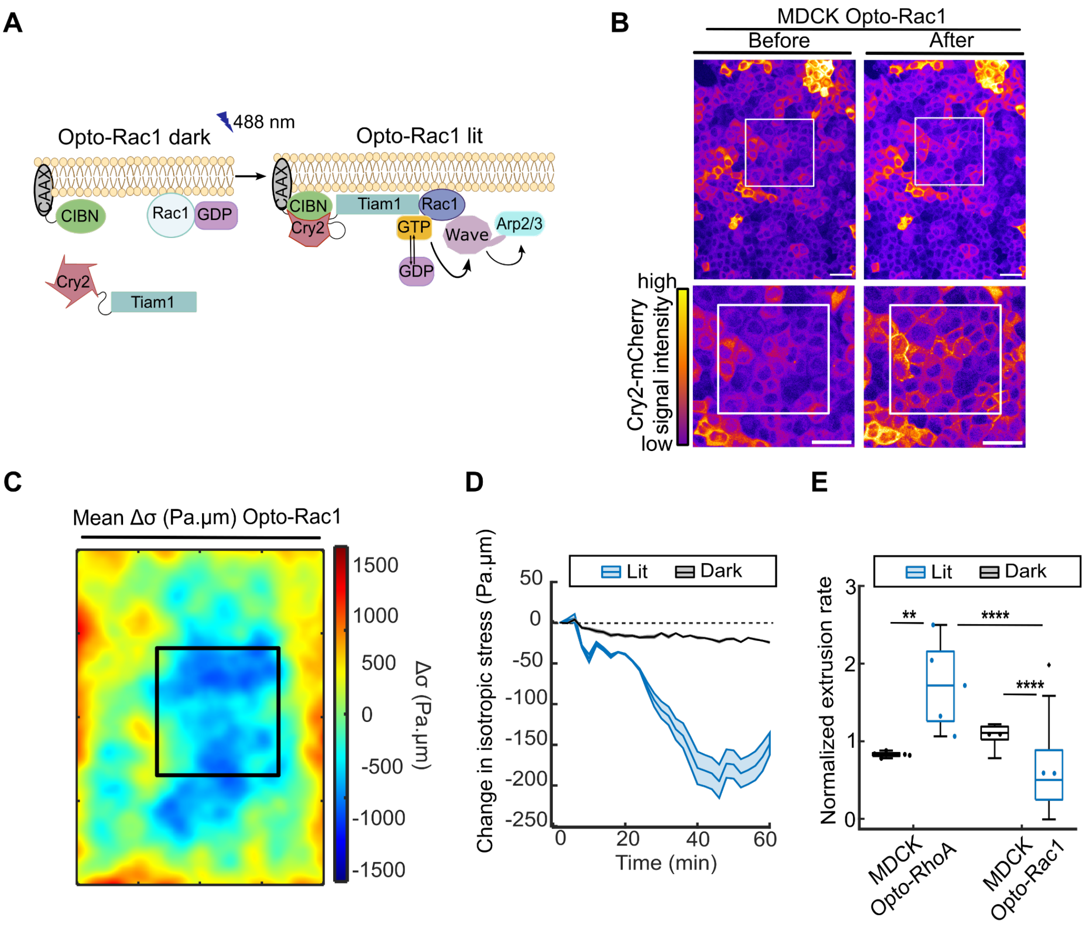


**Figure S6: Opto-Rac1 induces a local decrease of isotropic stress and cell extrusions. Related to Figure 2.**

**A** Schematic of the optogenetic system used to trigger Rac1 activation in MDCK cells. (left) CRY2-Tiam1-mCherry is recruited to the plasma membrane following blue light-stimulated binding to CIBN, leading to Rac1 activation and subsequent Arp2/3 activation (Right). **B** Examples of membrane recruitment of Tiam1-CRY2-mCherry in MDCK Opto-Rac1 monolayer within a 150 x 150 µm square stimulated with 488nm blue light (white outline). Bottom: inlet magnification. **C** Map of change in isotropic stress of MDCK Opto-Rac1 monolayer 60 minutes after light activation in the 150 x 150 µm square stimulated with 488 nm blue light (white outline) (N=2 n=38). **D** Quantification of change in isotropic stress in illuminated regions for MDCK Opto-Rac1 (blue) or surrounding unstimulated regions (black) from mean maps shown in **(C)**. **E** Quantification of mean extrusion rate in the stimulated (lit) or non-stimulated (dark) regions normalized by their area sizes, for MDCK Opto-RhoA monolayers (N=5 n=51) and MDCK Opto-Rac1 monolayers (N=2 n=38). Wilcoxon rank-sum test: ** p = 0.007937; ** p = 0.001499; **** p = $2.72 \times 10^{-7}$, from top left to bottom right. Curves represent the mean +/- s.e.m. Box plots represent the median, interquartile (box), and 1.5 IQR (whiskers).

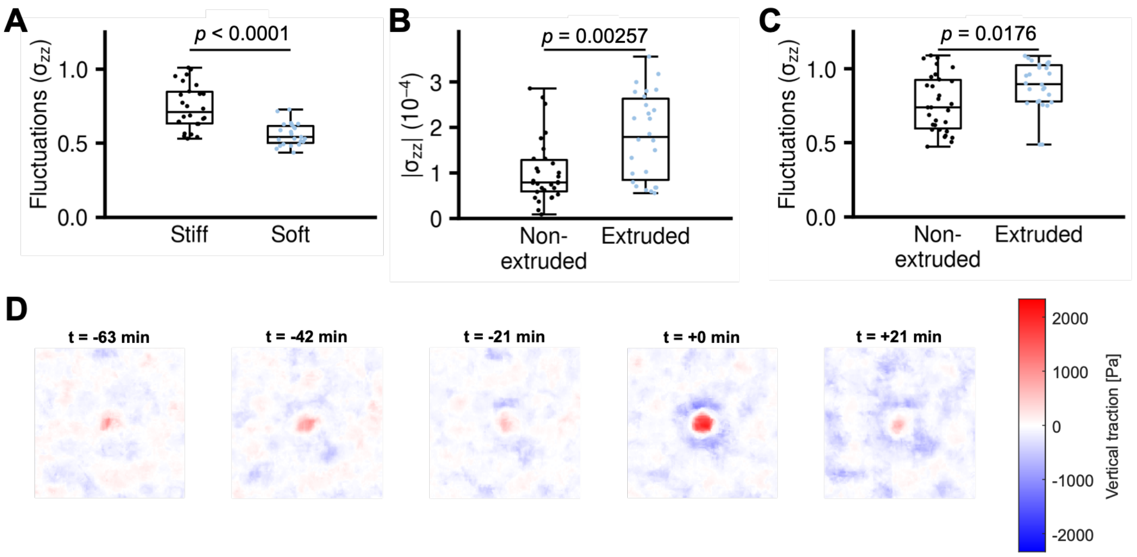


**Figure S7: Extrusions happen in regions of high fluctuations of out-of-plane stress in stiff cells. Related to Figure 3.**

**A** (From simulations) Quantification of mean $\sigma_{zz}$ fluctuations within the extrusion region confirms that extrusions in the stiff (activated) domain occur under significantly higher vertical fluctuations than those in soft regions (Stiff n = 26 events; Soft n = 23 events, from 3 independent simulations). **B, C** (From simulation) Within the activated domain, cells that extrude exhibit significant higher out-of-plane stress magnitude (B) and fluctuations (C, extruded n = 26 events, non-extruded n = 30 events, from 3 independent simulations). Mann–Whitney test; Box plots represent the median (center line), interquartile range (box), and minimum to maximum values (whiskers). **D** (From experiments) Vertical traction forces were measured from 2.5 D TFM experiments in 40 x 40 µm regions centered on extrusion events and averaged over multiple positions (N=2 n=38, see Figure 3I).

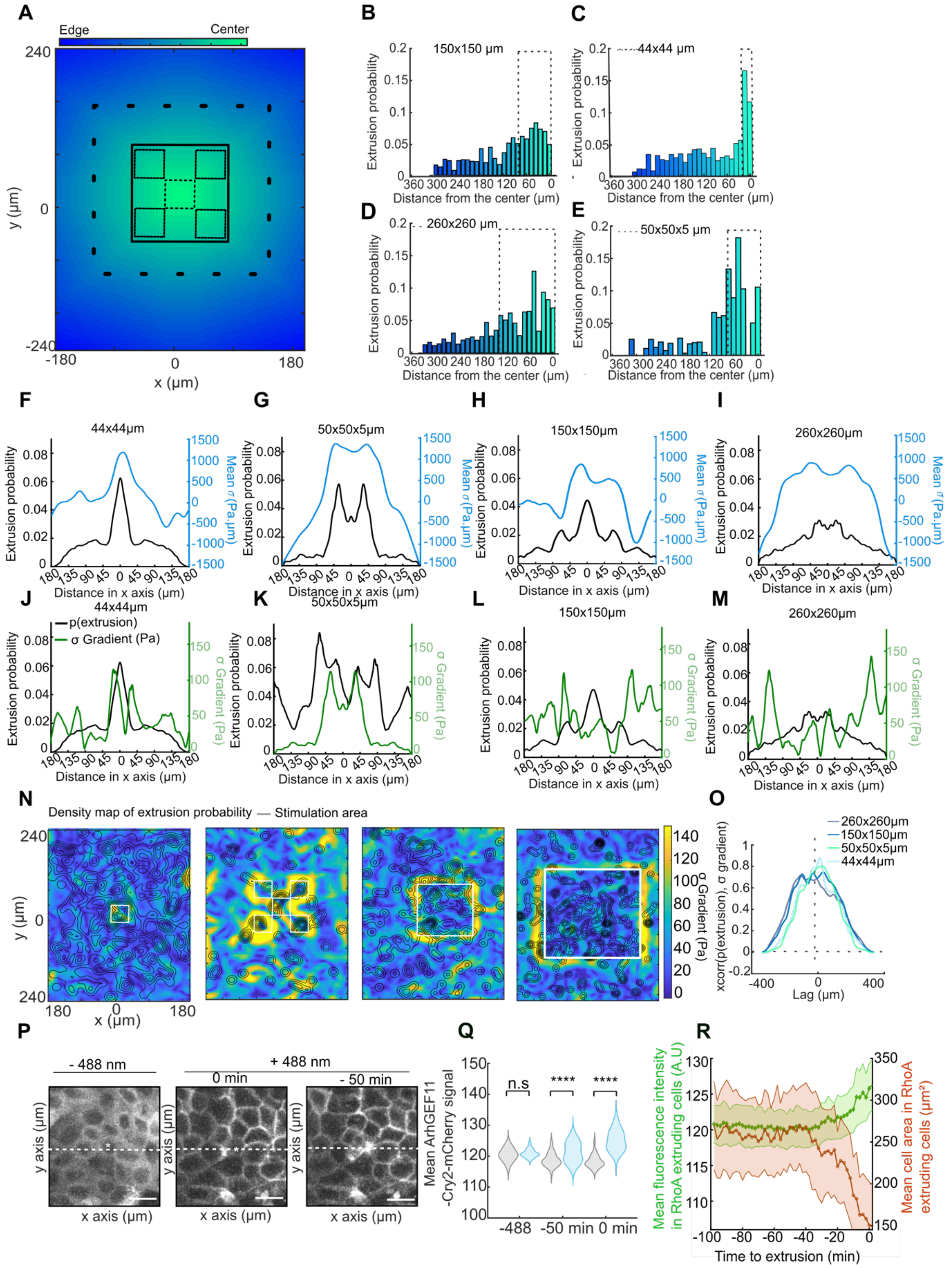

A
Edge
Center
y (µm)
x (µm)
B
150x150 µm
C
44x44 µm
D
260x260 µm
E
50x50x5 µm
Extrusion probability
Distance from the center (µm)
F
44x44µm
G
50x50x5µm
H
150x150µm
I
260x260µm
Mean σ (Pa.µm)
Distance in x axis (µm)
J
44x44µm
p(extrusion)
σ Gradient (Pa)
K
50x50x5µm
L
150x150µm
M
260x260µm
σ Gradient (Pa)
N
Density map of extrusion probability
Stimulation area
σ Gradient (Pa)
O
260x260µm
150x150µm
50x50x5µm
44x44µm
xcorr(p(extrusion), σ gradient)
Lag (µm)
P
- 488 nm
+ 488 nm
0 min
- 50 min
y axis (µm)
x axis (µm)
Q
Mean ArhGEF11 -Cry2-mCherry signal
n.s
****
****
-488
-50 min
0 min
R
Mean fluorescence intensity in RhoA extruding cells (A.U)
Mean cell area in RhoA extruding cells (µm²)
Time to extrusion (min)

**Figure S8: Extrusion probability correlates with stress amplitude rather than stress gradient. Related to Figure 4.**

**A** Scheme describing the distance maps of brightfield images used to quantify extrusion probability for different size of stimulation patterns: 260 x 260 µm (green), 150 x 150 µm (blue), 44 x 44 µm (pink), and 50 x 50 chessboard regions (red), as presented in Figure 4A. **B,C,D,E** Distribution of extrusion probability as a function of the distance from the center. (N=3 n=47, N=5 n=51, N=2 n=32, N=2 n=32 positions, respectively). Stimulated areas are represented by dark dashed rectangles. **F,G,H,I** Spatial distribution of extrusion probability and mean isotropic stress as a function of distance. **J,K,L,M** Spatial distribution of extrusion probability and magnitude of isotropic stress gradient as a function of distance. **N** Density maps of extrusion locations (black level curves) and magnitude of isotropic stress gradient (color coded) in the increasing order of regions stimulation size (from left to right, white squares). **O** Normalized cross-correlation functions between the spatial distribution of cell extrusion probability and magnitude of isotropic stress gradient in these different conditions. **P** Representative fluorescence images of Cry2mCherry signal in MDCK Opto-RhoA monolayers before stimulation (−488 nm) and during blue light stimulation (+488 nm), shown 50 minutes before extrusion onset (−50 min) and at extrusion onset (0 min). Dashed white lines indicate the regions used for fluorescence intensity profile measurements. **Q** Distribution of ArhGEF11–Cry2-mCherry fluorescence intensity in MDCK Opto-RhoA cells before blue-light stimulation (−488 nm), 50 min prior to extrusion onset (−50 min), and at the onset of extrusion (0 min). Violin plots were generated from the average fluorescence intensity measured within a 10 × 10 µm region centered on the future extrusion site (blue) and within surrounding cells (gray) (N=4, n=20). Wilcoxon rank-sum test: n.s., $p = 0.1548$; **** $p = 3.59 \times 10^{-6}$; **** $p = 4.02 \times 10^{-21}$, from left to right. **R** Curves of mean Cry2-mCherry fluorescence intensity (green) and mean cell area (orange) in stimulated extruding Opto-RhoA cells (N=4, n=20) as a function of time relative to extrusion onset (t = 0 min). Scale bars: 10 µm (magnified insets).

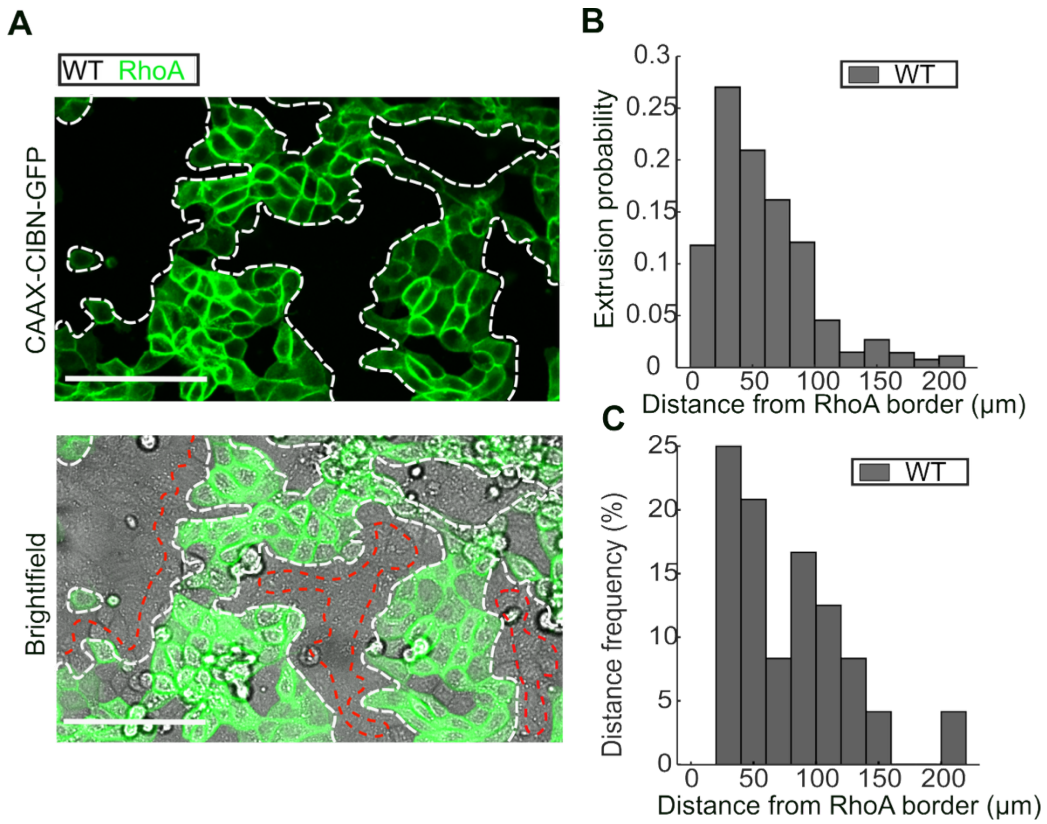


**Figure S9**: **Extrusion of WT cells does not depend on the distance with opto-RhoA cells. Related to Figure 4.**

**A** Representative image of co-culture experiments composed of 50% MDCK Opto-RhoA cells and 50% wild-type MDCK cells. MDCK Opto-RhoA cells are identified by CAAX-GFP labeling (top). The white dashed line denotes the border between the RhoA and WT populations, and the red dashed line indicates a distance equivalent to one WT cell row from the interface. **B** Distribution of extrusion probability normalized of MDCK WT cells as a function of the distance from the MDCK Opto-RhoA border (N=3, n=148 cells). **C** Distribution of the frequency of distances between MDCK WT cells and the border of MDCK Opto-RhoA cells, showing that MDCK WT cells are most frequently located at less than 50 µm of the MDCK Opto-RhoA border.

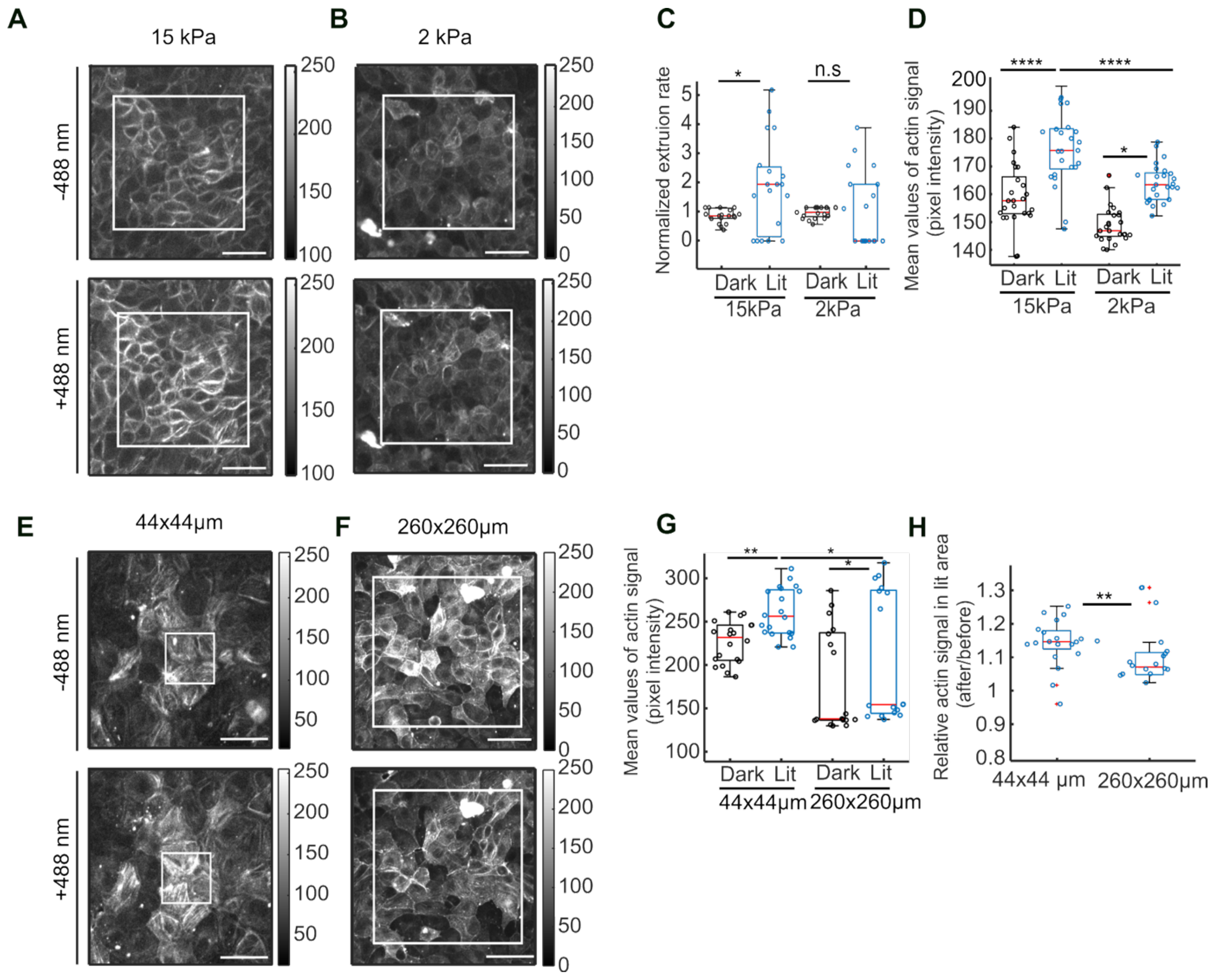


**Figure S10: Enrichment of stress fibers are required for tension mediated extrusions. Related to Figure 5.**

**A,B** Actin labeling in Opto-RhoA cells cultivated on 15 kPa (A) and 2 kPA PDMS (B), before (top) and after (bottom) stimulation in a 150 x 150 µm square. **C** Normalized extrusion rate in the stimulated (lit) or non-stimulated (dark) regions in MDCK Opto-RhoA monolayers cultured on 15 kPa (N=2 n=20) and 2 kPa (N=2 n=20). Wilcoxon rank-sum test: * p = 0.04042; n.s., p = 0.6091, from left to right. **D** Mean actin signal at the basal side in the stimulated (lit) or non-stimulated (dark) regions for MDCK Opto-RhoA monolayers cultured on 15k Pa (N=2 n=25) and 2 kPa (N=2 n=20). Wilcoxon rank-sum test: **** p = $6.41 \times 10^{-5}$; *** p = $1.04 \times 10^{-4}$; * p = 0.01794, from top to bottom and left to right. **E,F** Representative images of MDCK Opto-RhoA monolayer cultured on 15 kPa PDMS and stimulated in a region of 44 x 44 µm. Mean actin intensity before (top) and after (bottom) optogenetic stimulation in a 44 x 44 µm (E) or 260 x 260 µm (F) region. **G** Mean actin signal at the basal side in the stimulated (lit) or non-stimulated (dark) regions for MDCK Opto-RhoA monolayers stimulated in a region of 44 x 44 µm or 260 x 260 µm. Wilcoxon rank-sum test: ** p = 0.001953; * p = 0.05040; * p = 0.02750, from left to right. **H** Relative values of actin signal (actin signal after/before stimulation) at the basal side of MDCK Opto-RhoA cells stimulated in 44 x 44 µm (N=2, n=20) or 260 x 260 µm areas (N=2, n=20). Wilcoxon rank-sum test: * p = 0.01142. Box plots represent the median, interquartile (box), and 1.5 IQR (whiskers).

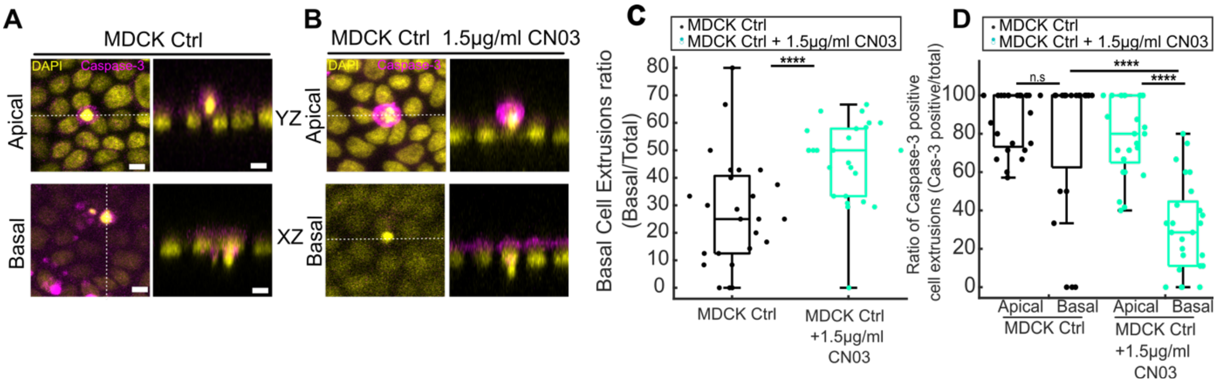


**Figure S11: RhoA activation by CN03 predominantly triggers live basal extrusions. Related to Figure 6.**

**A, B** Representative images of apical (top) and basal (bottom) extrusions of MDCK Opto-Ctrl cells grown on collagen I matrices, either untreated **(A)** or treated with 1.5 µg/ml CN03 Rho Activator (**B)**. Magenta: active caspase-3, yellow: DAPI. Side views of cell extrusions are shown on the right (white dashed lines, YZ and XZ planes). **C** Quantification of the ratio of basal cell extrusions (basal/total) in untreated (dark, N=2 n=207 cells) and treated conditions (cyan, N=2 n=370 cells). Wilcoxon rank-sum test: **** $p = 7.01 \times 10^{-5}$. **D** Ratio of caspase-3–positive extrusions (caspase-3–positive/total) for apical and basal extrusions in both conditions. Wilcoxon rank-sum test: n.s., $p = 0.4728$; **** $p = 3.23 \times 10^{-5}$;n.s., $p = 0.4728$ **** $p = 2.51 \times 10^{-7}$, from left to right. Scale bars: 10 µm (magnified insets). Box plots represent the median, interquartile (box), and 1.5 IQR (whiskers).

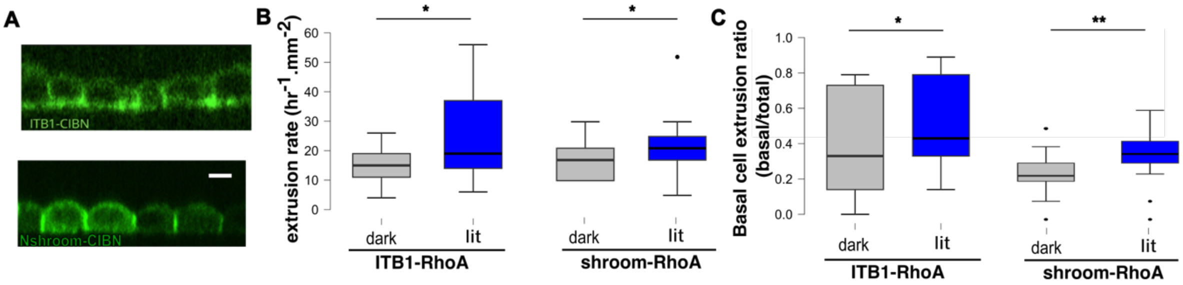


**Figure S12: Apical *vs.* basal activation of RhoA don't influence extrusion orientation/ related to Figure 6.**

**A** Confocal images of the GFP signal showing a baso-lateral localisation in ITB1-mVenus-CIBN;ArhGEF11-CRY2-mCherry cells (ITB1-RhoA, top) and an apico-lateral localisation in N-Shroom-CIBN-GFP;ArhGEF11-CRY2-mCherry cells (Shroom-RhoA, bottom). Scale bar: 10 µm. **B** Quantification of the extrusion rate in ITB1-RhoA (left) and Shroom-RhoA (right) cells, stimulated (lit, blue) or non-stimulated (dark, grey). N = 4, Wilcoxon rank-sum test: * p = 0.038; * p = 0.043 from left to right. **C** Ratio of basal/total extrusions in these conditions. Wilcoxon rank-sum test: * p = 0.041; ** p = 0.0062.

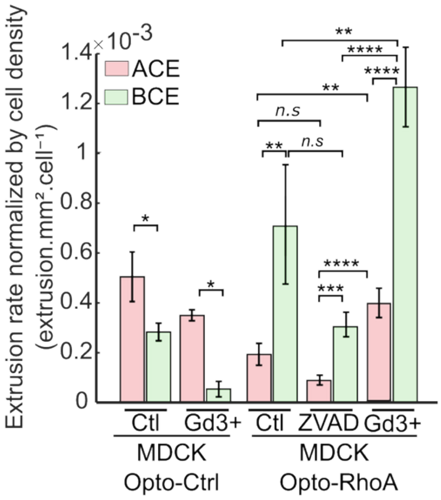


**Figure S13: Cell extrusion rate normalized to cell density under pharmacological inhibition of mechanosensitive channels and apoptosis. Related to figure 6.**

Quantification of the cell extrusion rate normalized by the cell density in illuminated MDCK Opto-Ctrl and MDCK Opto-RhoA monolayers under control conditions (Ctl) or following treatment with 50 µM gadolinium chloride (Gd3+) or Z-VAD-FMK (N=2 n=35, N=1 n=16, N=3 n=90, N=2 n=61, N=2 n=60, N=2 n=191 cells respectively). Bar plots represent the mean ± s.e.m. Wilcoxon rank-sum test: ** $p = 0.002138$; **** $p = 6.25 \times 10^{-6}$; **** $p = 2.67 \times 10^{-5}$;** $p = 0.003477$; n.s., $p = 0.07534$; ** $p = 0.001499$; n.s., $p = 0.06338$; * $p = 0.02857$; **** $p = 6.25 \times 10^{-6}$; * $p = 0.01499$; *** $p = 7.35 \times 10^{-4}$; from top to bottom.
